# Phages are important unrecognized players in the ecology of the oral pathogen *Porphyromonas gingivalis*

**DOI:** 10.1101/2022.12.30.519816

**Authors:** Cole B. Matrishin, Elaine M. Haase, Floyd E. Dewhirst, Jessica L. Mark Welch, Fabiola Miranda-Sanchez, Donald C. MacFarland, Kathryn M. Kauffman

## Abstract

**Background:** *Porphyromonas gingivalis* (hereafter “*Pg*”) is an oral pathogen that can act as a keystone driver of inflammation and periodontal disease. Although *Pg* is most readily recovered from individuals with actively progressing periodontal disease, healthy individuals and those with stable non-progressing disease are also colonized by *Pg*. Insights into the factors shaping the striking strain-level variation in *Pg*, and its variable associations with disease, are needed to achieve a more mechanistic understanding of periodontal disease and its progression. A key force shaping strain level diversity in all microbial communities is infection of bacteria by their viral (phage) predators and symbionts. Surprisingly, although *Pg* has been the subject of study for over 40 years, essentially nothing is known of its phages, and the prevailing paradigm is that phages are not important in the ecology of *Pg*.

**Results:** Here we systematically addressed the question of whether *Pg* are infected by phages - and we found that they are. We found that prophages are common in *Pg*, they are genomically diverse, and they encode genes that have the potential to alter *Pg* physiology and interactions. We found that phages represent unrecognized targets of the prevalent CRISPR-Cas defense systems in *Pg*, and that *Pg* strains encode numerous additional mechanistically diverse candidate anti-phage defense systems. We also found that phages and candidate anti-phage defense system elements together are major contributors to strain level diversity and the species pangenome of this oral pathogen. Finally, we demonstrate that prophages harbored by a model *Pg* strain are active in culture, producing extracellular viral particles in broth cultures.

**Discussion:** This work definitively establishes that phages are a major unrecognized force shaping the ecology and intraspecies strain-level diversity of the well-studied oral pathogen *Pg*. The foundational phage sequence datasets and model systems that we establish here add to the rich context of all that is already known about *Pg*, and point to numerous avenues of future inquiry that promise to shed new light on fundamental features of phage impacts on human health and disease broadly.

## Introduction

One of the most actively studied and thoroughly described microbial ecosystems is that of the human mouth^1^. A major insight that has emerged from studies of the oral microbiome is that microbially-mediated oral inflammation is associated with increased risk for inflammatory disease throughout the body. Understanding the mechanisms underlying the most common oral inflammatory disease, periodontitis (periodontal disease), therefore has relevance to human health not only in the mouth but systemically.

*Porphyromonas gingivalis* (hereafter *“Pg”*) is a keystone pathogen in oral inflammation^2^, driving conditions that favor periodontitis. This gram-negative asaccharolytic anaerobe is adapted for growth in the gingival crevice, colonizes microbial biofilms through attachments to specific bacteria, and its growth is facilitated by growth factors produced by the community^3^. Once in the gingival crevice, *Pg* has the capacity to produce copious amounts of proteolytic enzymes that facilitate immune evasion and supply its preferred peptide nutrient source. Importantly, *Pg’s* proteolytic activity also disrupts the attachment of the junctional epithelium to the tooth surface, promoting formation of “pockets” of exposed epithelium around the teeth^4^, creating niches that favor the growth of other oral pathogens, and accelerating disease progression. Notably however, although *Pg* is most readily recovered from individuals with actively progressing periodontal pockets, healthy individuals and those with stable non-progressing pockets are also colonized by *Pg*. Identifying the factors that shape strain-level variation in the physiology, interactions, and virulence potential of *Pg* is thus an important element of achieving a comprehensive view of the role of this species in oral inflammation and systemic disease.

A key force shaping strain-level diversity in all microbial communities is infection of bacteria by their viral predators and symbionts, called bacteriophages (hereafter “phages”). Phages are highly specific in their infections, in part due to their need to bind to specific host bacterial surface moieties (e.g. lipopolysaccharide, capsule, or outer membrane proteins^5^) in a lock-and-key fashion. As a result, infection and killing by phages can exert a strong diversifying effect on microbial populations through negative frequency dependent selection (favoring rarer strains and genes). In the oral microbiome phages have been shown to be numerous and diverse, present at up to 10^8^ ml^-1^ in saliva and 10^10^ g^-1^ dental plaque^6^. Metagenomic studies of oral viromes have revealed the presence of phage-encoded virulence factors^7^, enrichment for genes predicted to shape bacterial interactions with human host cells^8^, and shifts in phage communities in periodontal disease^9^. Elegant early laboratory studies of the processes underlying oral biofilm community assembly also have revealed that the receptors used by oral phages include the very cell surface molecules key to co-evolved co-aggregation between different bacterial species (e.g. *Actinomyces* and *Streptococcus sp*.)^10–13^. This latter work suggests that negative selection pressure exerted by phages is an integral factor shaping dynamics of oral biofilm development *in vivo*. However, despite the potential for studies of phage-bacteria interactions to shed light on the ecology of specific bacteria and the structure and function of microbial communities, cultivated bacteria-phage model systems are lacking for all but a few species in the oral microbiome^14^.

The extent to which the key oral pathogen *Pg* interacts with phages remains a major open question. Surprisingly, although *Pg* has been the subject of study for over 40 years^15^, essentially nothing is known of its phages, and the prevailing paradigm is that phages are not important in the ecology of *Pg*. An early study^16^ using a culture-based approach to uncover prophage interactions in *Pg* found no evidence of plaque formation (lysis of bacteria in lawns), and though future studies with modified approaches were recommended there is no evidence they were carried out. More recent studies using comparative genomic analyses of *Pg* genomes have identified candidate phage genes^17^, and two *Pg* genomes have been noted as having candidate prophage regions, though these are not further described (ATCC 49417, with a region noted as Bacteriophage phi Pg1 in GenBank record FUFH01000018.1; and WW2952, with a candidate prophage noted as PgSL1^18^). Investigation of the targets of the highly prevalent CRISPR-Cas defense systems in *Pg* have also identified candidate phage genes in *Pg* genomes as potential targets^19^. Yet, because it is thought that there are no phages infecting *Pg^19^*, studies of these systems have highlighted other roles^20^, finding them to be independently associated with virulence^21^ and highly expressed in periodontal disease^22^. The presence of CRISPR-Cas systems in *Pg* genomes has been suggested to explain the lack of phages infecting this species^19^. Yet, the existence and prevalence of defense systems also implies that selection to maintain these may be due to predation pressure by as-yet-unrecognized phages infecting *Pg*.

Here, we sought to systematically address the question of whether *Pg* are, or are not, infected by phages - and we found that they are. Using a bioinformatic approach, we showed that integrated phages (“prophages”) are common in *Pg*, they are genomically diverse, and they encode genes that have the potential to alter their host physiology and interactions. We found that phages represent unrecognized targets of the prevalent CRISPR-Cas defense systems in *Pg*, and that *Pg* strains encode numerous additional mechanistically diverse candidate anti-phage defense systems. We also showed that phages and candidate anti-phage defense system elements together are major contributors to strain level diversity and the species pangenome of this oral pathogen. Finally, we find that nuclease-protected phage genomes and virus-like particles can be found in culture supernatants of a *Pg* strain encoding a prophage. In sum, this work reveals that interactions with phages are a major unrecognized force shaping the ecology and intraspecies strain-level diversity of the well-studied oral pathogen *Pg*.

## Results

### *Pg* isolates harbor phylogenetically diverse prophages

To address the question of whether *Pg* interacts with phages we focused our investigation on temperate phages, which have the capacity to integrate into their host bacterial genomes and form stable associations as prophages. Phages with temperate life history strategies are known to be common in the oral microbiome^23^ and offer the possibility of discovery based on study of bacterial genome sequences alone.

Definitive identification of prophages in bacterial genomes remains a challenge for the field. To systematically search for prophages in *Pg* genomes, we therefore used an approach that combined multiple complementary lines of evidence (Supplementary Fig. 1, and see Methods for details). In brief, we first analyzed the species pangenome to identify variable genomic regions not present in all *Pgs* and thus likely to include mobile elements such as phages. Next, we applied a panel of well-developed prophage prediction and annotation tools to identify the subset of variable genome regions likely to be prophages. We then harvested all CRISPR-spacers from identified arrays in *Pg*, as well as other species of bacteria, and mapped these back to all *Pg* genomes to facilitate detection of regions that are likely to be actively mobilizing. Finally, we identified precise boundaries of prophage regions by manual curation using all-by-all BLAST-based genomic comparisons among all *Pg* genomes, all available phage and bacterial annotations, and likely insertion site sequences and bounding repeats. Together these methods yielded comprehensive and high-quality prophage predictions.

Using our integrative approach, we found that prophages are common in strains of *Pg*, present in 32% (25/79) of strains examined (Fig. 1). We searched for prophages in all publicly available *Pg* genomes, as well as in four additional genomes we sequenced for this work (a total of 79 strains, hereafter “*Pg*_set_79”, and 88 genomes including cases of sub-strains and re-sequenced strains, hereafter “*Pg*_set_88”; see Supplementary Data 1). Four of the 25 *Pg* strains with prophages encoded two prophages each, and an additional four *Pg* harbored partial prophage regions. We also identified an additional *Pg* with a prophage likely incomplete only due to an assembly artifact (phage033a). The distribution of prophages with respect to the *Pg* phylogeny did not reveal obvious patterns of association with specific clades, nor suggest host-ranges dependent on use of receptors linked with virulence (see Supplementary Data 2), though a limited exploratory analysis did show an apparent bloom of phage in a periodontal disease metagenome (Supplementary Fig. 2). Determining the bacterial surface receptors used by these phages to infect *Pg* will be the subject of future studies. Defining these receptors is expected to provide new insight on selective pressures acting on *Pg* expression of cell surface moieties (e.g. O- and K-antigens, fimbriae, and other outer membrane proteins^5^) commonly used by phages to infect their hosts, and that play a role in virulence and capacity for *Pg* to bind to partners in oral biofilms (e.g. *Streptococcus gordonii*), recruit other species (e.g *Fusobacterium nucleatum*), bind to and invade human host cells, and avoid phagocytosis.

**Figure 1.**
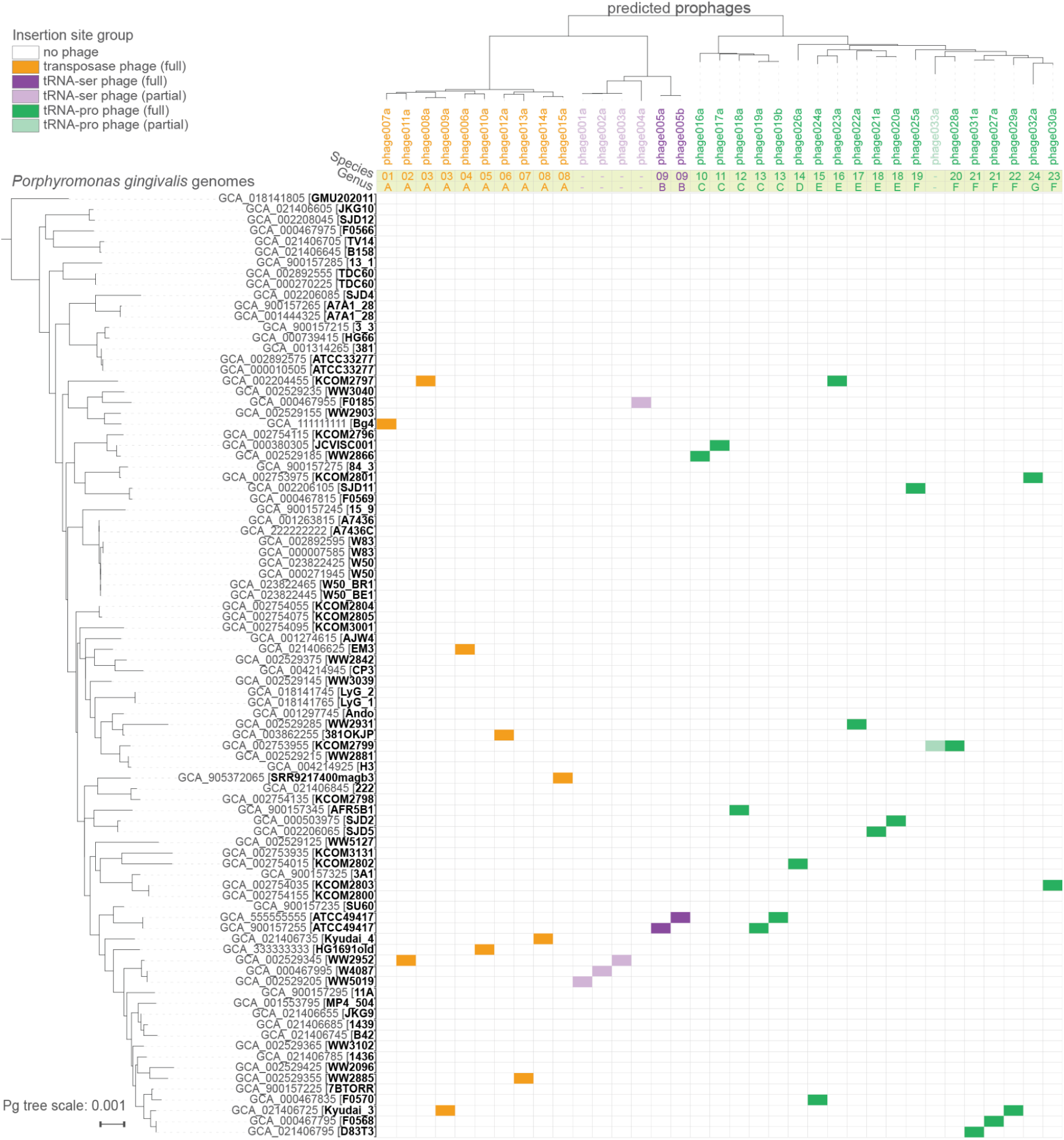
Prophages are common in sequenced *Porphyromonas gingivalis* isolates. Phylogenetic relationships among *Pg* shown on the left (79 strains; 88 leaves, including 3 substrains and 6 strains with independent assemblies), based on concatenated ribosomal protein genes. Relationships among *Pg* phages shown in midpoint-rooted tree at the top (30 full, 5 partial; “b” suffix indicates version of an “a” phage found in a different assembly of the *Pg* strain), based on whole genome BLAST distance (using concatenated proteins and scaled by VICTOR^74^ d6 formula). Candidate genus- and species-level clusters are shown for full-length phages in the yellow bars. Three higher order clades of phages defined by distinct insertion sites in host genomes (by full-length phages only) are highlighted (see color legend). Colored cells in the matrix indicate the assemblies in which each phage was found.

To examine the diversity and novelty of these predicted phages we compared them to each other and to non-*Pg* phage genome sequences (see Methods). The phages we found in *Pg* represent 7 new candidate genus-level groups comprising 24 species-level subgroups (Fig. 1). The overall diversity of these phages could be organized into three higher order clades defined by their use of distinct conserved insertion sites in *Pg* host genomes. Clade 1 was characterized by non-site-specific transposition-based insertion [present in 10 *Pg* strains], Clade 2 by insertion into tRNA-Pro [in 5 *Pg* strains], and Clade 3 by insertion into tRNA-Ser [in 17 *Pg* strains]). Genome sequence conservation was highest among phages sharing the same insertion site type (Supplementary Fig. 3).

### *Pg* phage genomes harbor genes with potential to shape host ecology

To understand the potential impacts of phages on *Pg* hosts that they infect we used numerous annotation databases and iterative HMM-based searches to predict the functions of their genes (see Methods, Supplementary Data 3). In general, the phage genes most readily annotated are those encoding structural and packaging components of the virion (e.g. capsid, tail, portal, terminase large subunit), and this held true for the *Pg* phages (Fig. 2). Based on sequence similarity and conservation of structural gene order, all phages identified here were predicted to be siphoviruses with long non-contractile tails. However, predicted structural and assembly genes together accounted for only 363 (19%) of the 1,892 genes in these 33 phages, and the majority of phage genes (56%) could not be readily annotated.

**Figure 2.**
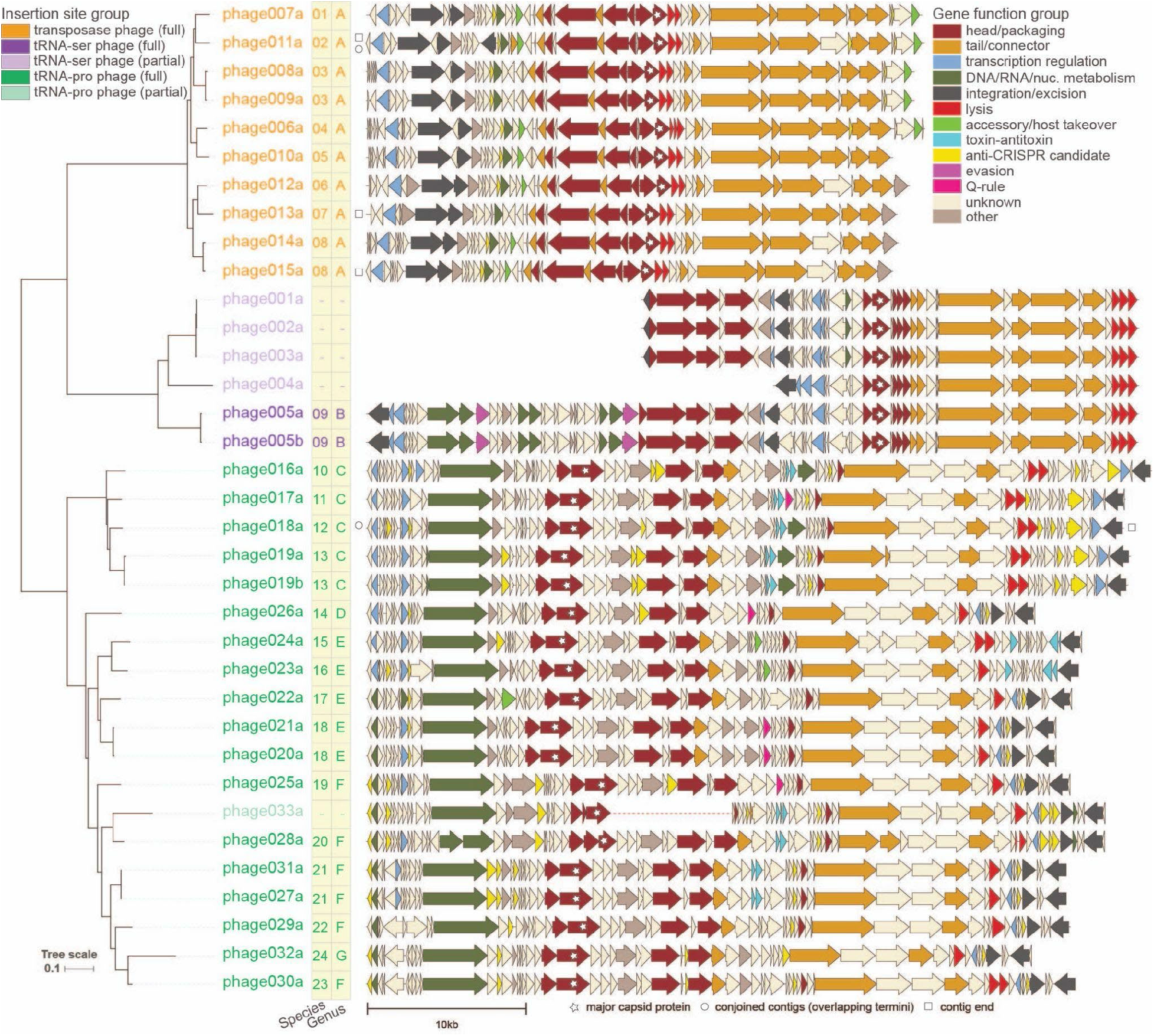
Genome diagrams of *Porphyromonas gingivalis* phages highlight functional annotations and gene order conservation in three large clades defined by distinct use of host genome insertion sites. Representations of *Pg* phage genomes (30 full, 5 partial; names of full length phages are in saturated colors and partial phages are in lighter shades; “b” suffix indicates version of an “a” phage found in a different assembly of the *Pg* strain), generated using Clinker^75^ and showing predicted protein coding genes as block arrows colored based on predicted protein functional categories (see Supplementary Fig. 3 for version with protein clustering). Relationships among *Pg* phages shown in midpoint-rooted tree at left, based on whole genome BLAST distance (using concatenated proteins and scaled by VICTOR^74^ d6 formula). Candidate genus- and species-level clusters are shown for full-length phages in the yellow bars. Three higher order clades of phages defined by distinct insertion sites in host genomes (by full-length phages only) are highlighted by coloring of phage names (orange: transposition based insertion; purple: tRNA-Serine; green: tRNA-Proline). White stars mark phage genome ends defined by contig ends, circles mark phage genomes identified in this work by joining contigs with overlapping termini, the dotted line in the middle of phage033a highlights that this phage was identified at the two termini of a bacterial contig assembly and is missing genes potentially due to an incomplete assembly.

Despite the challenges of annotating phage genes, several intriguing classes of genes with the potential to impact *Pg* physiology and virulence emerged in this first investigation. Among these were the putative lipopolysaccharide (LPS)-modifying enzymes, proteins with signal peptides targeting them to transport by the general secretion system, and toxin-antitoxin systems; we highlight these examples below.

First, the majority of the transposable *Pg* phages encode putative phosphoheptose isomerases, genes that participate in LPS synthesis (light green genes in orange phage group in Fig. 2). The presence of LPS-modifying genes in phages has previously been shown to result in modifications of host bacterial LPS that alter bacterial virulence potential and sensitivity to infection by related phages^24,25^. That this gene is common in the transposable phage clade suggests that LPS is a receptor for this group, as it is for the related transposable phage Mu^26^. This finding points to transposable phages having the potential to alter *Pg* ecology and virulence not only through inactivation of genes upon non-specific integration into bacterial genomes, but also through modification of host LPS, a key contributor to *Pg* virulence.

Second, multiple phages in the tRNA-Pro clade encode genes with signal peptide sequences. A subset of these genes are associated with core phage functions, including major capsid proteins whose signal peptides are likely cleaved by the phage encoded prohead proteases, and spanins, which are necessary for lysis. Yet strikingly, five genes of unknown function with signal peptides are predicted to obey the *“Bacteroidetes* Q Rule” of revealing exposed N-terminal glutamine residues once processed^27^ (neon pink genes in green phage group in Fig. 2). That *Pg* phages encode proteins that follow the Q-rule, a unique and distinctive feature of Signal Peptidase I substrates in the *Bacteroidetes^27^*, suggests that they are adapted to using their host’s general secretion systems and have the potential to modify *Pg* outer membranes and thereby their interactions.

Third, phages in the tRNA-Pro clade also commonly encode toxin-antitoxin (TA) system genes (neon blue genes in green phage group in Fig. 2, Supplementary Data 3). TA systems are mechanistically diverse but share the property of encoding a toxin that reduces bacterial metabolic activity and an anti-toxin that neutralizes the toxin. These systems are upregulated in bacteria as defenses in response to phage infection^28,29^, and as a survival strategy during other cellular stress events; for example, TA systems can induce a persister state upon exposure to antibiotics or nutrient starvation^30^. Although TA systems encoded on phages may play a role in promoting maintenance of these selfish elements in their host populations, they have also recently been shown to act in inter-phage competition^31^, preventing successful infections of the host by other phages. The diverse roles of TAs in bacterial physiology raise the question of whether *Pg* prophages encoding TAs can provide an ecological advantage to their hosts in the stressful subgingival crevice^32^. The most readily recognizable TA systems in the *Pg* phages are Type II HicAB dyads that function by degrading mRNA, reversibly reducing global translation^32,33^. Additional singleton toxins and antitoxins are also present in the prophages. Solo antitoxin genes encoded on phage genomes have been shown to act as counter-defenses to bacterial TA-mediated attempts to abort infections^34^. Solo toxins, however, are not expected, and as we found these in genomic islands, known to be used by phages to harbor anti-phage genes^31^, we expect that partner genes for these singletons will ultimately be identified among nearby genes. Future studies examining expression of integrated prophage TA genes in *Pg* strains across physiologically relevant growth conditions (including exposure to predation by exogenous phages) are needed to reveal whether they play a role in promoting *Pg* survival.

### Prophages are targets of *Pg* CRISPR systems and encode putative anti-CRISPR genes

Finding that prophages are common in *Pg* raised the question of whether they represent targets of spacers in *Pg* CRISPR arrays. *Pg* strains commonly encode CRISPR arrays, yet the targets of the spacers have remained elusive^19,35^. To address the question of potential phage targeting by *Pg* CRISPR systems, we harvested the spacers from *Pg* genomes in our dataset using CRISPRCasTyper^36^ (CCTyper) and compared these with the sequences of the prophages (Fig. 3, see Methods).

**Figure 3.**
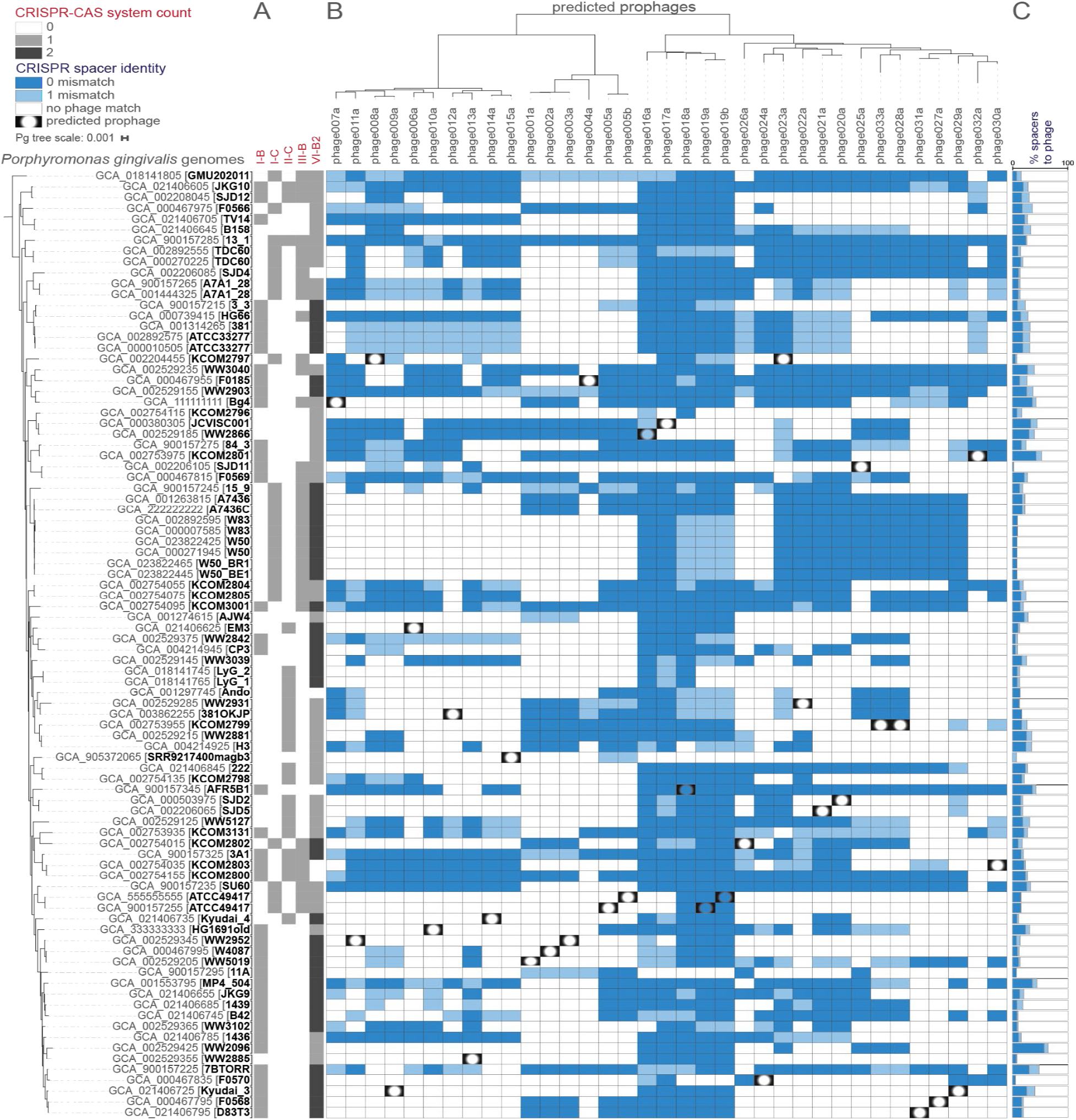
*Porphyromonas gingivalis* CRISPR arrays encode spacers that target phages in other strains. Predicted CRISPR-Cas systems in each strain of *Pg* are shown; quantities of each system are related to cell color saturation (A). CRISPR spacer hits from arrays found in *Pg* are mapped onto *Pg* phages shown in midpoint-rooted tree at the top (30 full, 5 partial; “b” suffix indicates version of an “a” phage found in a different assembly of the *Pg* strain; based on whole genome BLAST distance (using concatenated proteins and scaled by VICTOR^74^ d6 formula); dark blue cells indicate 0-mismatch spacer-phage nucleotide identity, light blue indicates 1-mismatch, and vignetting indicates presence of the entire prophage in the bacteria (as shown in Fig. 1) (B). Percent of total spacers found in each *Pg* that have 0- or 1-mismatch to a predicted phage are shown; same coloring as panel B (C). CRISPR-Cas systems were identified by CCTyper^36^ and mapped to phage genomes with Bowtie^76^. Phylogenetic relationships among *Pg* are shown on the left (79 strains; 88 leaves, including 3 substrains and 6 strains with independent assemblies), based on concatenated ribosomal protein genes.

Overall, we found that prophages account for a substantial fraction of targets of CRISPR array spacers in *Pg*. Every *Pg* strain we investigated encodes at least one CRISPR-Cas system (Fig. 3A), and their arrays collectively encode 4,993 spacers (*Pg*_set79, see Supplementary Data 4). Considering spacers in all *Pg* genomes, a total of 833 (17%) showed 100% nucleotide identity to at least one of the phages characterized in this study (Fig. 3B), and 1150 (23%) mapped to phages if allowing 1 nucleotide mismatch in the alignment (see Supplementary Data 5). Considering spacers in individual strains of *Pg*, we found that the proportion targeting phages can be far larger, with up to 57% of spacers in a given strain being sequence-identical to phages in this collection (up to 64% if allowing 1 mismatch in mapping, Fig. 3C). Given that we expect that the diversity of *Pg* phages exceeds that which we have captured in our study in this limited number of genomes, we also considered the possibility that unaccounted for spacers might target phages more distantly related to those in our dataset (e.g. showing protein conservation but nucleotide divergence). To address this we mapped six-frame spacer translations against *Pg* phage-derived peptides using SpacePHARER^37^ and found that this increased the proportion of spacers that could be accounted for by phages to 27% (1,342/4,993, Supplementary Data 6).

As expected, the majority of *Pg* strains do not carry spacers that map to their own prophages, yet we noticed that a small number do (3 of 28 strains with prophages in *Pg*_set79; see filled-in “peepholes” in Fig. 3B). In two cases there are only few matching spacers, however, strain AFR5B1 appears to extensively target its own phage018a (14 0-mismatch hits from a Class 1 Type I-B array, and 3 more if allowing 1 mismatch, see Supplementary Data 5). The large number of matches to phage018a in AFR5B1 was striking and raised the question of whether this phage encodes anti-CRISPR protein genes (phage counter-defense genes protecting against CRISPR-Cas systems), that would have allowed it to survive targeting upon infection to successfully achieve integration^38^.

To investigate whether genes encoding anti-CRISPRs (acrs) are present *Pg* phages we used tools and databases designed for their discovery, including PaCRISPR^39^ and the DeepAcr database^40^. To reduce candidates to those of highest confidence we considered only those predicted to be acrs by both PaCRISPR and DeepAcr, and found that these represented 26 phage protein coding gene families, encompassing a total of 99 proteins (see Supplementary Data 3). Mapping the candidate acrs to phage genomes we found that they occur in variable regions enriched in small, often hypothetical, genes (Fig. 2, yellow genes). In the genome of phage018a, mentioned above as being heavily targeted by spacers in its own parent *Pg* genome, we found six candidate acrs. Though these predicted acrs require future study for validation, our findings suggest that such genes may indeed have played a role in the successful integration of phage018A into AFR5B1 by inhibiting CRISPR-Cas targeting^38^. *Pg* prophages thus offer plentiful candidate acrs for future *in vitro* functional validation and characterization of phage genes involved in the bacteria-phage arms race in the human oral microbiome.

We also took advantage of CRISPR spacers to explore whether these *Pg* phages may have hosts in other bacterial species. To do this, we used CRISPROpenDb^41^, which maps >1.3 million unique spacers harvested from CRISPR arrays in 1,978 bacterial genera to potential targets. We found that matches between array spacers in *Porphyromonas gulae* (a species found in dogs, and the nearest phylogenetic neighbor to *Pg*) and *Pg* prophages were common (242 100% identity matches, see Supplementary Data 7). This suggests that phages infecting both of these closely-related bacterial species are likewise closely-related and may have the potential to recombine upon co-infection, and in exploratory analyses we found that *P. gulae* harbored predicted prophages in the same genera as the *Pg* phages described here.

In characterizing *Pg* CRISPR-Cas systems we found a striking near-ubiquity of Class 2 Type VI systems, generally rare among bacteria^42^. Although Type VI-B are the most common system in *Pg*, the majority of spacers were encoded by Type I-B rather than Type VI-B arrays (Type I-B: 51%, I-C:17%, II-C: 11%, III-B: 4%, VI-B2: 17%, Supplementary Data 4). Despite their relatively lower abundance, we found Type VI array spacers targeting all 24 candidate species of *Pg* phages at 100% identity. Recent work has demonstrated that Type VI systems offer broad spectrum activity against phages^42^, likely in part due to their lack of a requirement for protospacer associated motifs, this suggests the possibility that fewer spacers are needed by these systems to achieve coverage of diverse phages. Of note, whereas Type VI effector genes were commonly identified (Cas13b, and the Cas13b-activated membrane pore-forming Csx28^43^, identified as tm_HEPN by CCTyper^36^), genes associated with the adaptation module (e.g. cas1 or cas2) were not. In some cases it has been shown that Type VI-B systems can acquire spacers in trans from other co-occurring systems (e.g. from Type II-C systems in *Flavobacterium^44^*), however, we found no spacers or repeats shared between Type VI-B and any other array types in *Pg* (Supplementary Fig. 4, Supplementary Data 4). This suggests that novel Type VI CRISPR-Cas adaptation modules will be identified among the conserved hypothetical genes near Type VI effector genes in future studies.

Overall, our finding that *Pg* CRISPR-Cas arrays are enriched for spacers that target phages confirms that, in addition to their potential roles in bacterial physiology and virulence^21^,^45^, defense against phage infection is one of their major functions in this species. The unceasing bacteria-phage arms race is reflected here in the numerous candidate acrs we found in phages. Recent work has highlighted the complexity^46^ and specificity^47,48^ of interactions between defense and counter defense systems and unraveling the structure of these interactions to predict phage host ranges at the bacterial strain level remains a major challenge for the field.

### Non-CRISPR-Cas defense systems are also common and diverse in *Pg* genomes

Given that differences in carriage of prophages and anti-phage defense systems are a major source of intra-species diversity in other bacteria, we asked to what extent this is true for *Pg*. To expand our investigation of defense beyond CRISPR-Cas systems, we screened all 88 *Pg* genomes for the presence of any of the >100 defense systems in the PADLOC-DB v1.4. We found that there are at least 31 non-CRISPR-Cas systems in *Pg* (Fig. 4, Supplementary Data 8), including abortive infections systems, restriction-modification systems, retron-based interference systems (e.g. Septu^49^), and systems that use cyclic nucleotides to activate effectors (e.g. CBASS^50^ and Thoeris^49^).

**Figure 4.**
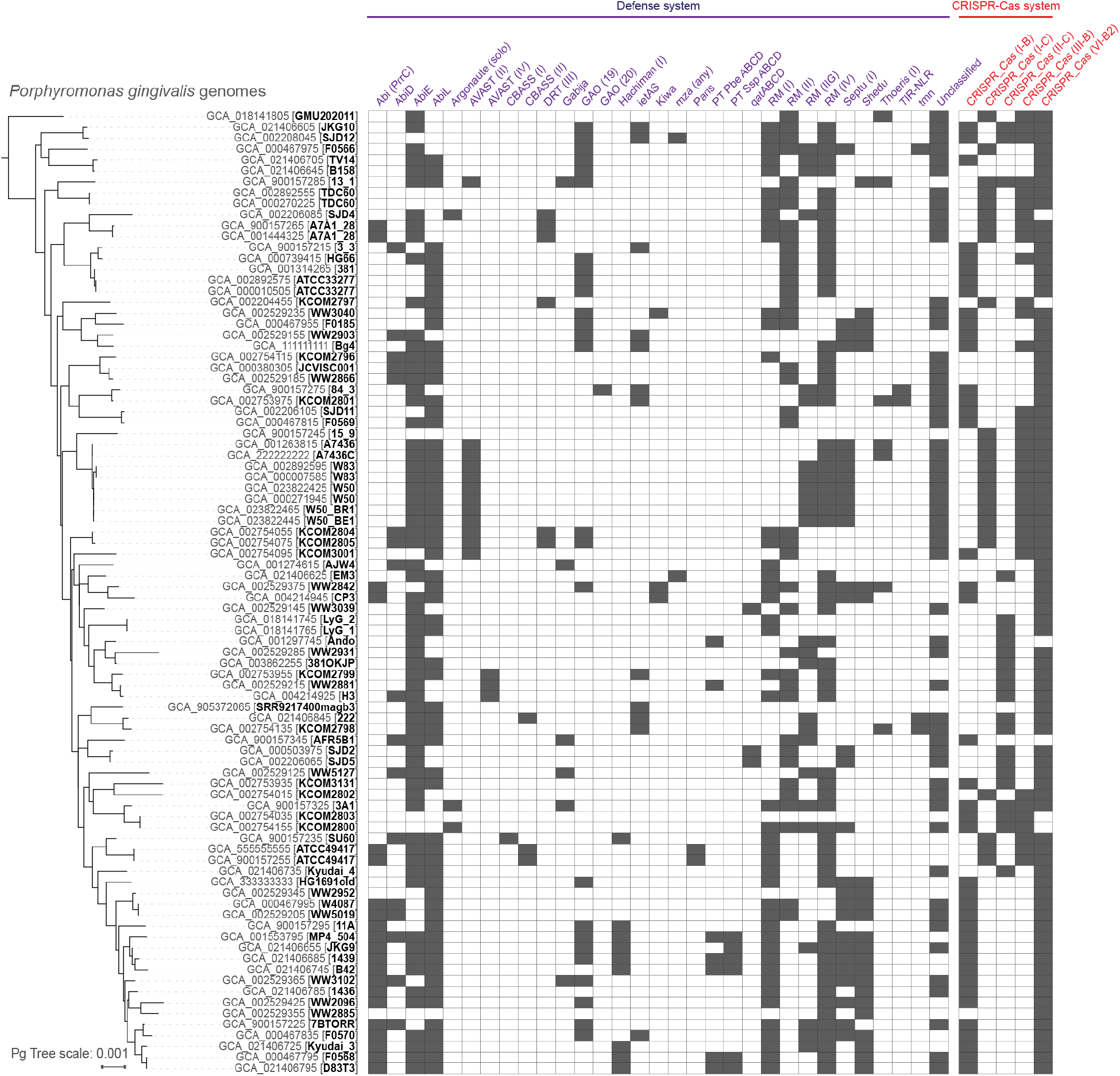
The *Porphyromonas gingivalis* species level pan-immune system is diverse. Presence of defense systems in each strain of *Pg* are indicated by filled in cells. Systems identified by PADLOC^55^ shown in panel A (excluding CRISPR-Cas systems) with subtypes indicated within parentheses where applicable, CRISPR-Cas systems identified by CCTyper^36^ shown in panel B. Phylogenetic relationships among *Pg* are shown on the left (79 strains; 88 leaves, including 3 substrains and 6 strains with independent assemblies), based on concatenated ribosomal protein genes.

As previously highlighted in the context of the bacteria-phage arms-race, “Where there is defense, there is counter-defense”^51^. One example of such a counter-defense system is an immune evasion associated nuclease (Anti-Pycsar, Apyc1) predicted with high confidence in phage005 (purple gene in Fig. 2, gene phage005a_ATCC49417_12 in Supplementary Data 3). Anti-Pycsar nucleases allow phages to escape bacterial immune systems by degrading the cyclic-nucleotides that activate defense effectors^52^. That we did not identify a pycsar system in these *Pgs* suggests either that they are present in strains outside this collection, or that anti-pycsar-like nucleases target additional classes of bacterial immune systems. Though in depth searches for phage counter defense or immune evasion genes are beyond the scope of this work, the finding of one such system points to the presence of others among the numerous hypothetical genes in *Pg* prophages and this example provides a valuable model system for further characterization of the phage-bacteria arms race in the oral microbiome.

### Prophages and defense-related islands are a major part of the *Pg* pangenome

We next used a pangenome-based analysis to evaluate the overall relative contributions of prophages and defense islands to the *Pg* pangenome. Using an approach that considered conservation of gene ordering rather than numeric prevalence thresholds alone (PPanGGOLiN^53^), we clustered all protein coding genes to identify genes present in nearly all *Pg* (“core” gene families; >87% of genomes in this dataset) and genes that are variably present in *Pg* genomes (“flexible” gene families) (see Methods and Supplementary Fig. 5). At the level of individual *Pg*, we found that almost a quarter (23%) of every strain’s genome is comprised of flexible genes not shared by all members of the species. Across all genomes together we found 5,745 gene families (*Pg*_set_79), with 1,476 (26%) core, and the remaining 4,269 (74%) flexible (Supplementary Table 1), with overall nearly half of all gene families (48%) of unknown function (based on Bakta^54^ annotations).

The curation of phages described in this work allowed us to identify 8% of flexible *Pg* pangenome protein clusters as encoded by prophages (351/4,269 gene families, Supplementary Table 1, Supplementary Data 8). To also obtain an estimate of the contribution of defense systems to the *Pg* pangenome, we quantified gene families associated with regions of genome plasticity (PPanGGOLiN “RGPs”, comprised of runs of adjacent flexible genes) encoding either CRISPR-Cas systems (as predicted by CCTyper, Fig. 4) or other defense systems (as predicted by PADLOC^55^ and by manual annotation, see Methods and Fig. 4). Using this approach we found that 38% of flexible protein clusters (1,636/4,269, Supplementary Table 1, Supplementary Data 8) were encoded on likely defense islands. Thus, prophages and putative defense elements together comprise 46% of flexible gene families in the *Pg* pangenome.

Given our systematic curation of prophages in this dataset, we expect our estimate of the relative contribution of prophages to the *Pg* pangenome to remain fairly stable as new *Pg* isolates are sequenced in the future. However, our estimate of the contribution of predicted defense islands to the *Pg* pangenome is likely to be conservative, as it relies on functional annotation of genes. Islands of genes related to defense against phages are known to be major contributors to strain level diversity in environmental bacteria^47^,^49^,^56^, and with further study numerous additional gene families of currently unknown function in *Pg* will likely be revealed as defense systems. Altogether, these findings demonstrate that prophages, and the defense systems that protect against them, are important contributors to strain-level diversity in *Pg*.

Although we focused on the phages and defense systems for this work, transposons were also notably prominent in the pangenome. In particular, Insertion Sequence (IS) transposases are highly diverse (represented by at least 3,448 genes in 61 protein clusters, see Supplementary Data 9) and abundant in *Pg*, with some genomes having as many as 117 transposases. These findings are consistent with previous studies that have established IS elements as highly abundant and diverse in *Pg^57^*, contributing to gene regulation^58^ and genome recombination and targeted by CRISPR systems^20^. New in this work is our observation that one IS element is also present on a transposable phage (phage011a). Sequence comparisons reveal that the IS element present on the phage has >98% sequence identity over its entire length to elements in *Pg* strains Bg4, KCOM2797, and Kyudai3, but not to any IS’s in its own host, WW2952. This case suggests the possibility that the IS elements so ubiquitous in *Pg* genomes may be hitching rides on phages, using them as vectors of horizontal gene transfer, and benefiting from the phage’s capacity for immune evasion and counter defense.

### Prophages in *Pg* are active in culture

Finally, to investigate whether there is evidence for activation of *Pg* prophages in culture, we conducted laboratory studies focusing on a model strain (ATCC 49417) predicted to encode two functional prophages. These studies revealed the presence of abundant extracellular, nuclease-protected phage DNA from one of the two phages in culture supernatants (Fig. 5). Aged broth cultures of ATCC 49417 were filtered through 0.2 μm filters to remove bacterial cells and ultracentrifuged at 174,900 x g to pellet cell-free particles. Ultracentrifuge pellets were nuclease-treated to remove unprotected DNA prior to extraction, and Illumina sequencing revealed a high coverage enrichment of DNA from the region of the predicted siphovirus phage005, as compared with sequence from the background bacterial chromosome (Fig. 5, Supplementary Fig. 6, Supplementary Data 10). Electron microscopy of material from the resuspended pellet showed highly abundant particles of irregular size (presumably outer membrane vesicles)(Fig. 6a) as well as phage-like particles (Fig. 6b) similar to those observed in previous exploratory imaging studies of the same strain directly from broth cultures (Fig. 6c,d). Together, these observations indicate that cultures of *Pg* encoding prophages can produce cell-free nuclease-protected phage DNA and virus-like particles under common laboratory conditions.

**Figure 5.**
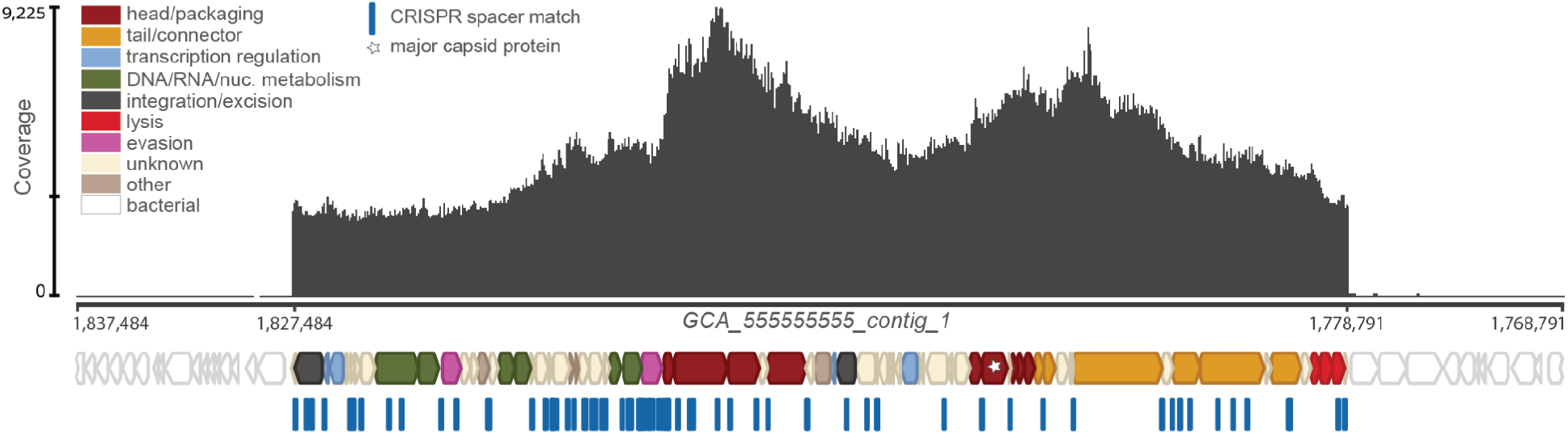
Protected phage DNA is present in *Pg* cultures. Coverage (dark gray plot) of nuclease-protected DNA sequences from a filtered 20-day old ATCC 49417 culture mapped onto a section of the ATCC 49417 genome that was assembled from the same untreated, 1-day old culture. An ~49 kb spike in coverage, with maximum 9,225x coverage (indicated by the scale on the left, middle hash mark notes mean coverage), corresponds with the region of phage005b in GCA_555555555_contig_1. Colored block arrows represent phage genes (major capsid protein marked with star), while white block arrows represent host genes predicted by Bakta^54^. CRISPR spacer matches to phage005b found in other *Pg*, predicted by CCTyper^36^ and mapped with Bowtie^76^ (100% identity), are represented by blue dash marks.

**Figure 6.**
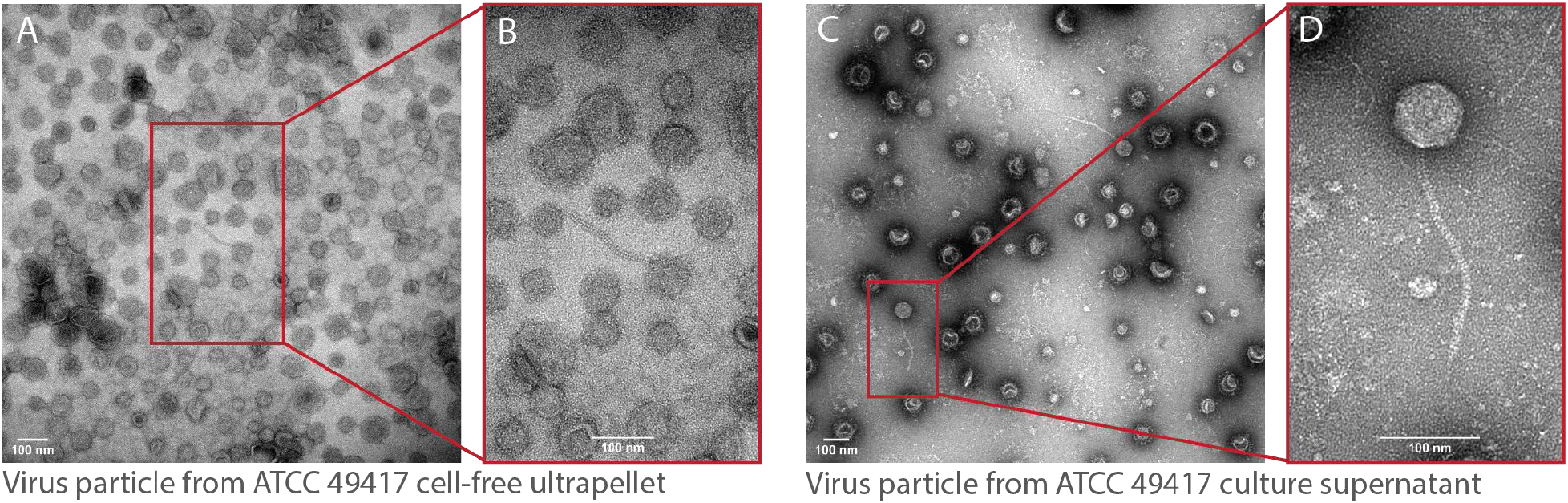
Virus-like particles are produced by *Pg* cultures. Transmission electron micrograph (2.73 pixels/nm) of the ATCC 49417 cell-free, ultracentrifuged supernatant that, when sequenced, produced the reads mapped in Fig. 5; virus-like particles were sparse among likely extracellular vesicles, despite high coverage of the prophage region in DNA from this material (**A**). Magnification (5.52 pixels/nm) of the virus-like particle from panel A (**B**). Transmission electron micrograph (2.25 pixels/nm) of supernatants of a 3-day aged ATCC 49417 culture derived from passages of the same stock that gave rise to the cultures imaged in panels A and B; virus-like particles were more abundant than in ultracentrifuge pellets shown in panels A and B and more commonly showed angular, icosahedral capsids (**C**). Magnification (8.81 pixels/nm) of the virus-like particle from panel C (**D**).

## Discussion

The discovery in this work that *Pg* are commonly infected by phages, and commit significant genomic real estate to predicted defense elements to protect against them, raises many new questions about this well-studied pathogen. Three broad areas of special interest for future studies are highlighted here.

### Dynamics of *Pg* prophage activation

In this work we showed that a strain of *Pg* harboring prophages naturally releases phage particles and DNA into the supernatant under standard broth culture conditions. Natural release of phages is known in other systems, including the related gut-associated *Bacteroides fragilis^59^*. However, we observed that in a *Pg* host with two prophages, only one phage dominated in the supernatant of broth cultures, and recent work has also shown that where bacterial strains harbor multiple prophages, these can have distinctive induction (activation) profiles^60^. This raises the question of what the natural cues are that different prophages are “listening in on” *in vivo* in the gingival crevice. Knowledge of which inducers are relevant in the oral microbiome has implications for understanding when there may be increased rates of cell lysis mediated by phage replication, and how different types of phages differentially impact lysis across conditions. Increased cell lysis has the potential to contribute to biofilm development through the release of free DNA, stimulate inflammation through release of bacterial cell debris, and facilitate horizontal gene transfer by multiple mechanisms. Previous studies have shown that a natural inducer in the oropharynx is hydrogen peroxide, which, for example, when produced by *Streptococcus pneumoniae* allows it to displace and outcompete *Staphylococcus aureus* competitors through “remote-control” induction of prophages^61^. Studies of the response of *Pg* ATCC 49417 (the strain for which we demonstrated phage release in culture) to hydrogen peroxide^62^ show limited effects on growth, suggesting that at least for the two groups of phages represented in our model hydrogen peroxide may not be the main inducer *in vivo* (though see below about non-canonical phage model systems).

Although the majority of phage genomes we identified are complete, and we have shown that they can form free phage particles, the copious production of vesicles by *Pg* also raises the question of whether phages can use vesicles as a mode of transmission. Packaging of phages into vesicles has been reported, and DNA packaged into vesicles by ATCC 49417 has been shown to be transferred and expressed in ATCC 33277^63^. Related, we also observed reproducible presence of nuclease-protected non-phage DNA in ATCC 49417 supernatants, suggesting that specific regions of *Pg* genomes are potentially packaged into vesicles and raising the question of whether this is associated with prophage activation. A recent study^64^ focusing on understanding dynamics of one of the most abundant groups of phages in the gut microbiome, the obligately lytic crAssphages infecting the *Bacteroides*, found that they do not form plaques in standard phage assays nor reduce turbidity of broth cultures of their hosts, though they are actively replicating. In general, it appears that phage infection dynamics in human-associated *Bacteroidetes* may commonly diverge from expectations based on studies of canonical model systems (e.g. *E. coli* infecting lambda- and T-phages). Resolving dynamics of prophage activation and spread in *Pg* model systems thus will likely also provide insight into phage-bacteria interactions in the human microbiome generally.

### Impacts of integrated phages on *Pg* physiology

In addition to identifying genes that likely alter *Pg* surface properties, we also showed that *Pg* phages harbor numerous genes of as-yet unknown function. A recent study^65^ demonstrated the power of transcriptomics to identify genes expressed by otherwise quiescent prophages, and revealed these as candidate modulators of host physiology across growth conditions (e.g. starvation, exposure to macrophages). Similar studies of new *Pg* phage model systems will be important for achieving a mechanistic understanding of how these phages are impacting *Pg*. In addition, the pressure for *Pg* strains to harbor defense systems against phages may impose fitness costs reflected in trade-offs between sensitivity to phages and growth rate. An elegant study^66^ in a marine system showed that the majority of bacteria in a population are resistant to phages yet are also slower growing compared with the rare phage-sensitive strains. To what extent the trade-off between defense and growth holds true for phage-bacteria interactions in the human microbiome is an important open question.

### Role of phages in oral colonization by *Pg* in health and disease

Since their discovery in the early 1900s, phages have been considered as offering potential to protect humans from disease caused by bacterial pathogens through their targeted killing^67^. More recently, phages in the human microbiome have been shown to bind to human mucins, forming a line of defense against colonization by pathogens^68^. Our finding that *Pg* strains commonly harbor prophages raises the question of whether phages also play a role in intra-species antagonism in this species in the mouth. Such dynamics would have the potential to limit colonization by new strains of *Pg*, either at the level of the person or at the level of individual periodontal pockets, as a result of killing by resident *Pg* phages. Recent work has also shown that in the related gut-associated *Bacteroides fragilis*, activated prophages are an important mechanism of intra-species antagonism and cross-killing^59^, and in Sandmeier’s search for *Pg* prophages in 1993^16^ he observed antagonism between strains of *Pg*, though he could not link it to phages. As individuals with *Pg* often harbor multiple strains of the species, with increasing numbers observed in periodontal disease^69^, the potential for phage-mediated intra-species antagonism also raises the possibility that bursts of disease progression are the result of bouts of phage-mediated cross-killing. Such a model was proposed in early studies of *Aggregatibacter* phages, where phage activity correlated with local disease progression^70^. Of note, if cross-kill dynamics are an important mechanism of periodontal disease progression, the presence or absence of specific phages alone is not expected to be predictive. Instead, the important property of the system will be the extent to which the specific strains of *Pg* colonizing an individual have the ability to antagonize one-another through phage-mediated mechanisms. That is, the specific structure of phage-bacteria interactions within the individual would matter for predicting outcomes. Understanding the determinants of host-range for *Pg* phages is thus an essential next step, and includes defining both the receptors used by phages and the relationships between the interacting bacterial defense and phage counter-defense (or immune evasion) genes.

Ultimately, understanding the roles of *Pg* phages in health and disease will also require broader sampling and study of clinical isolates, metagenomes, transcriptomes, and “live” phages from oral samples. The recent update of the largest metagenomic phage database (IMG/VRv4^71^), released in the late stages of our investigation, revealed numerous predicted *Pg* phages in *Pg* genomes and oral clinical metagenomic samples. Exploratory analyses of this new dataset showed that the majority of phages we identified and characterized here are represented in IMG/VRv4, though as the products of automated predictions their boundaries are less precise and not manually curated, as ours are. That several populations of predicted *Pg* phages present in IMG/VRv4 are not in our dataset underscores that their diversity exceeds what is represented in current *Pg* strain collections, and may also include exclusively lytic phages not discoverable in bacterial genomes. Worth remembering in all studies is that just as *Pg* exerts an outsized impact relative to its abundance, *Pg* phages are also exerting their effects from within a milieu of far more abundant phages. Judgements on the presence and activity of *Pg* phages must therefore be made in the context of datasets that are expected to have enough sequencing depth to detect them. Given these constraints, primers targeted to conserved regions of *Pg* phage genomes may provide a useful approach for initial screening studies aiming to broadly assess prevalence and associations of specific phages across states of health and disease.

## Conclusion

This work establishes that phages are important in the ecology of the oral pathogen *Pg*. The foundational phage sequence datasets and model systems that we establish here add to the rich context of all that is already known about *Pg* and point to numerous new avenues of inquiry. Given the challenges of understanding the complexities of phage-bacteria-human interactions, new model systems in the uniquely well-characterized^1,72,73^ context of the oral microbiome promise to shed new light on fundamental features of phage impacts on human health and disease broadly.

## Supporting information

Supplementary_Data_1.OVERVIEWINFOS_v2022.364

Supplementary_Data_2.VIRULENCE_v2022.364

Supplementary_Data_3.PHAGEANNOS_v2022.364

Supplementary_Data_4.CCTYPERANNOS_v2022.364

Supplementary_Data_5.CCTYPERHITSPHAGES_v2022.364

Supplementary_Data_6.SPACEPHARER_v2022.364

Supplementary_Data_7.CODBHITS_v2022.364

Supplementary_Data_8.BAXANNOS_v2022.364

Supplementary_Data_9.TRANSPOSASES_v2022.364

Supplementary_Data_10.ULTRAPEAKS_v2022.364

---

**Supplementary Figure 1.**
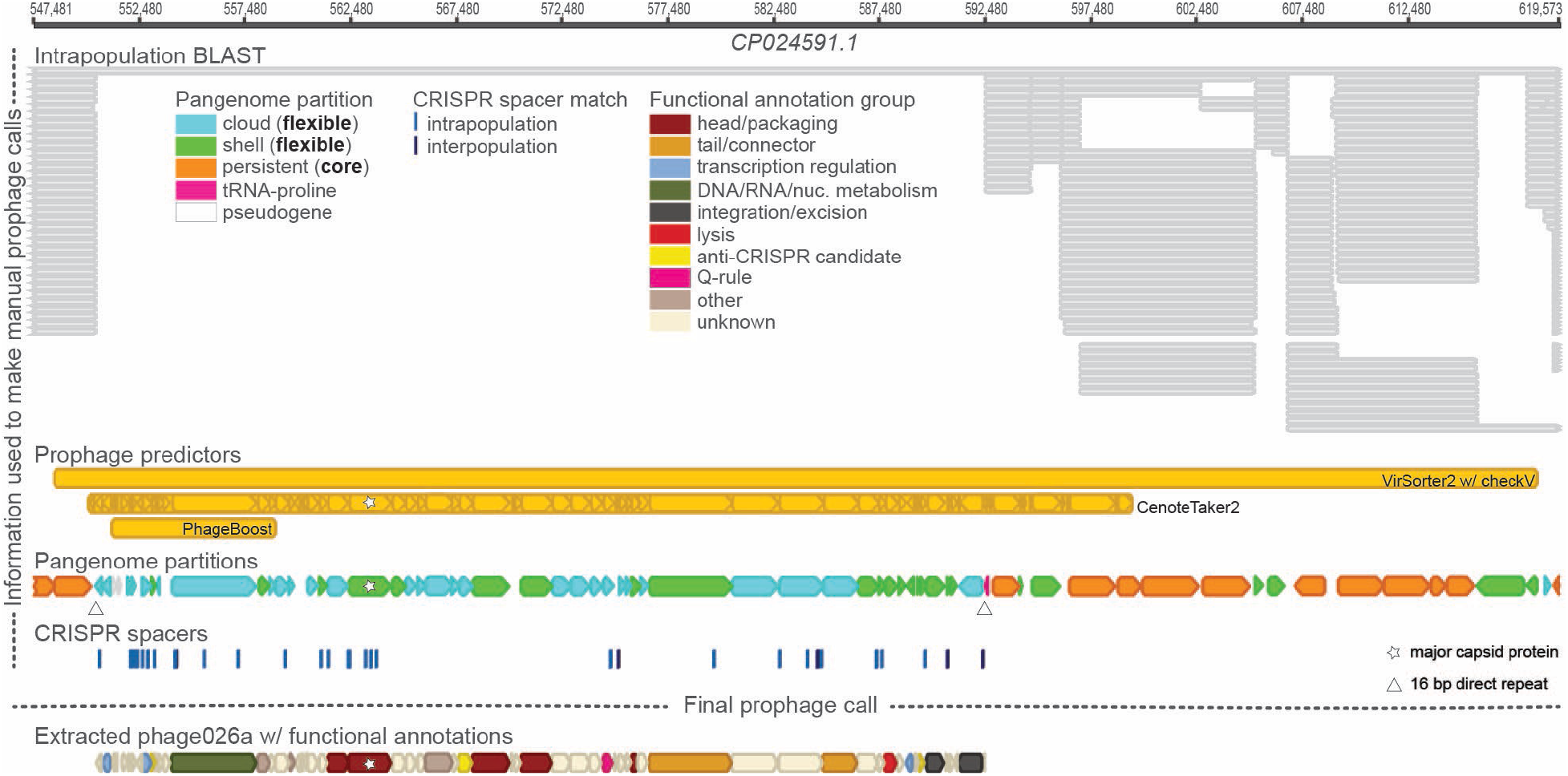
Rigorous bioinformatic approaches unveiled numerous *Pg* prophages. Example view from Geneious bioinformatic software highlighting numerous analyses utilized in manually curating *Pg* prophages. Bacterial contig CP024591 (KCOM 2802) was searched with five prophage predicting tools (VirSorter2^77^ with CheckV^78^, Cenote-Taker2^79^, PhageBoost^80^, VIBRANT^81^, and Inovirus Detector^82^); hits indicated in yellow bars. Annotations performed by Cenote-Taker2^79^ aided in determining the validity of the phage predictions due to powerful detection of major capsid proteins (marked by white stars). Pangenome partitions, predicted by PPanGGOLiN^53^, designate “flexible” protein-coding genes (light blue and light green block arrows), as compared to those that are “core” (orange block arrows); direct repeats were also identified as an indicator of insertion (those used by the phage marked by white triangles). CRISPR spacer matches (100% identity) found within other *Pg* strains (shown as blue hash marks; identified by CCTyper^36^) and other species (shown as dark blue hash marks; mapped from CRISPROpenDB^41^) elucidate regions targeted by intra- and interpopulation CRISPR-Cas systems, respectively. All-by-all intrapopulation BLAST used to compare each *Pg* genome against all other *Pg* genomes shows areas that lack conservation; hits indicated by gray bars. Prophage region (phage026 with functional annotations), inserted into a tRNA-Proline (pink block arrow) is manually curated, taking into account all the performed analyses.

**Supplementary Figure 2.**
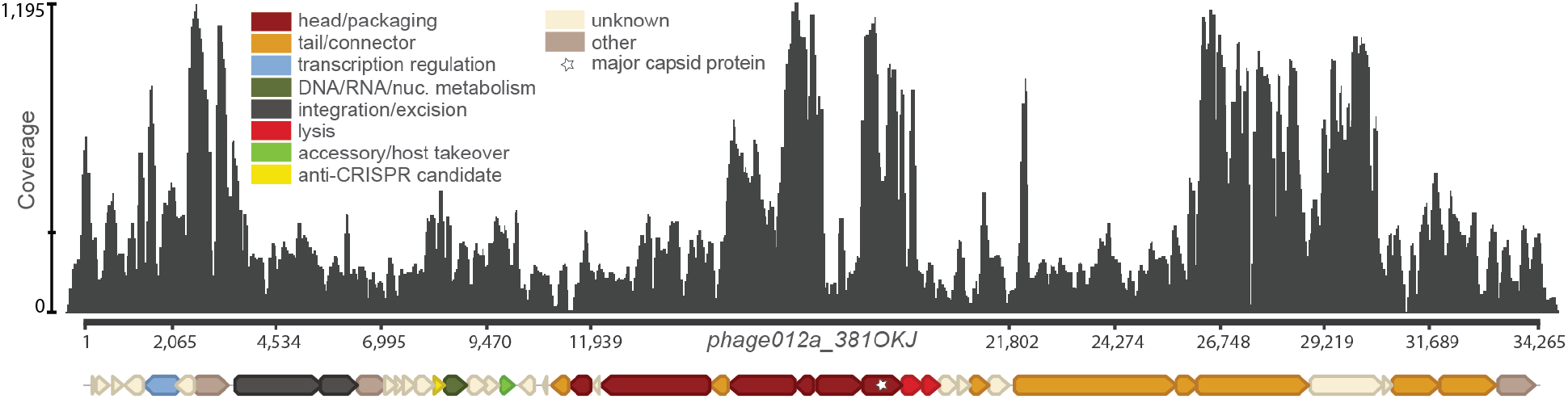
Transposable *Pg* phages are enriched in metagenomic reads from a periodontitis patient. Coverage (dark gray plot) of a transposable phage (phage012a_381OKJ) by metagenomic sequences sampled from the oral cavity of a periodontitis patient. Reads mapped to the entire phage genome, with maximum 1,195x coverage (indicated by the scale on the left, middle hash mark notes mean coverage). A preliminary search with these reads sequenced from the same patient showed lower coverage mappings to other transposable *Pg* phages.Colored block arrows represent phage functional gene groups (major capsid protein marked with star) predicted by Cenote-Taker2^79^.

**Supplementary Figure 3.**
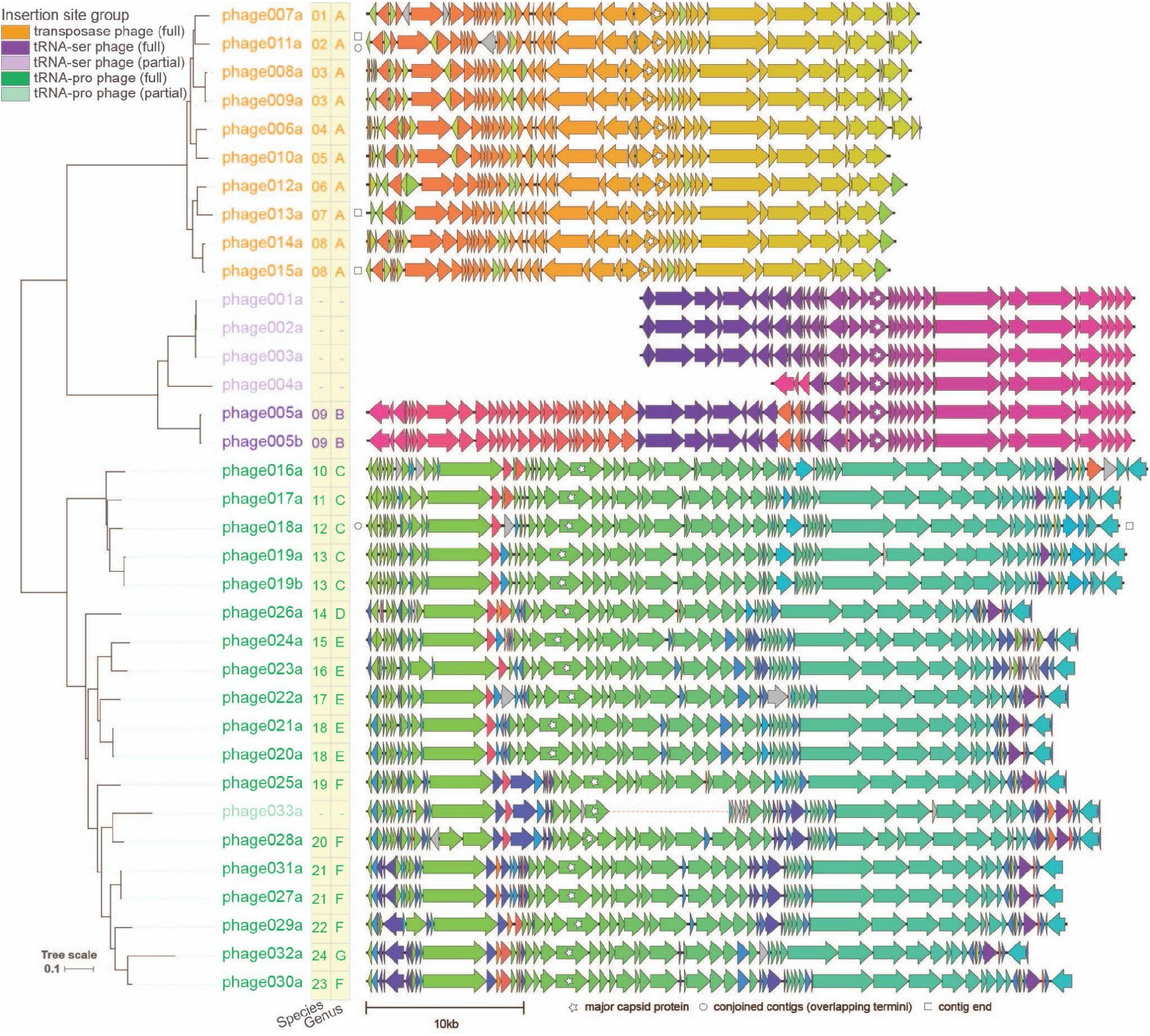
Genome diagrams of *Porphyromonas gingivalis* phages show conservation of protein clusters. *Pg* phage phylogeny (30 full, 5 partial; names of full length phages are in saturated colors and partial phages are in lighter shades; “b” suffix indicates version of an “a” phage found in a different assembly of the *Pg* strain; midpoint-rooted tree based on whole genome BLAST distance using concatenated proteins and scaled by VICTOR^74^ d6 formula) and genome diagram (generated using Clinker^75^) as shown in Fig. 2, with the exception that the predicted protein coding genes (depicted as block arrows) are colored based on sequence similarity. Thus, highlighting the conservation of protein clusters and ordering among related *Pg* phages. Candidate genus- and species-level clusters are shown for full-length phages in the yellow bars. Three higher order clades of phages defined by distinct insertion sites in host genomes (by full-length phages only) are highlighted by coloring of phage names (orange: transposition based insertion; purple: tRNA-Serine; green: tRNA-Proline). White stars mark phage genome ends defined by contig ends, circles mark phage genomes identified in this work by joining contigs with overlapping termini, the dotted line in the middle of phage033a highlights that this phage was identified at the two termini of a bacterial contig assembly and is missing genes potentially due to an incomplete assembly.

**Supplementary Figure 4.**
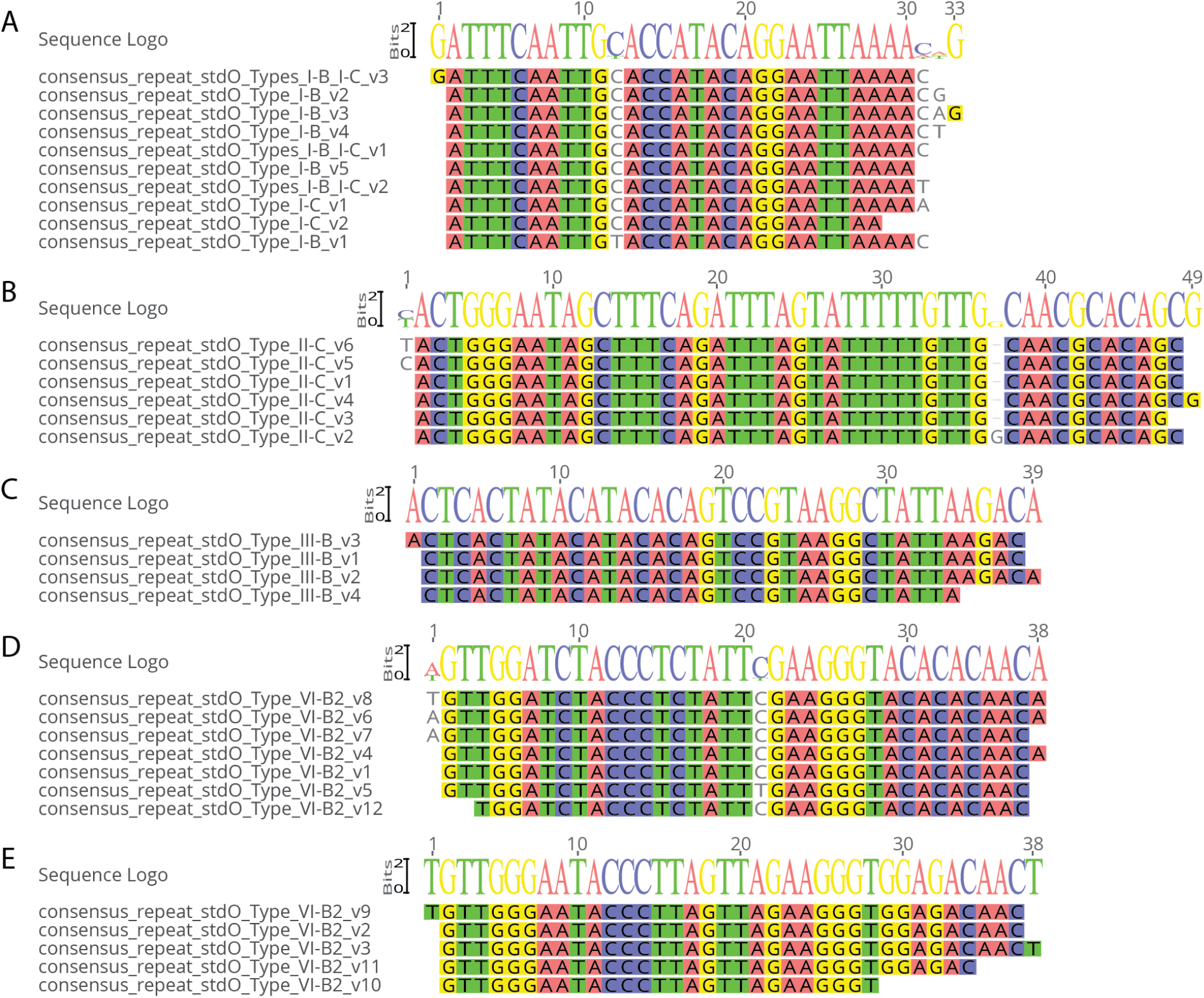
Consensus sequences for repeats within individual CRISPR-Cas arrays show intermingling within some systems and sequence diversity within others. Type I-B and I-C arrays share common consensus repeats (A), whereas Type II-C (B) and Type III-B arrays have distinct consensus repeats. Notably, for the Type VI-B arrays, there are two distinct groups of conserved repeats, one of these groups is associated with Type VI-B arrays that are part of the *Pg* core genome (D), whereas the other group (E) is associated with flexible Type VI-B systems. Consensus repeats shown are those from *Pg*_set79, in standardized orientation, and excluding repeats for which no CRISPR-Cas system type could be predicted (underlying data available in Supplementary Data 4).

**Supplementary Figure 5.**
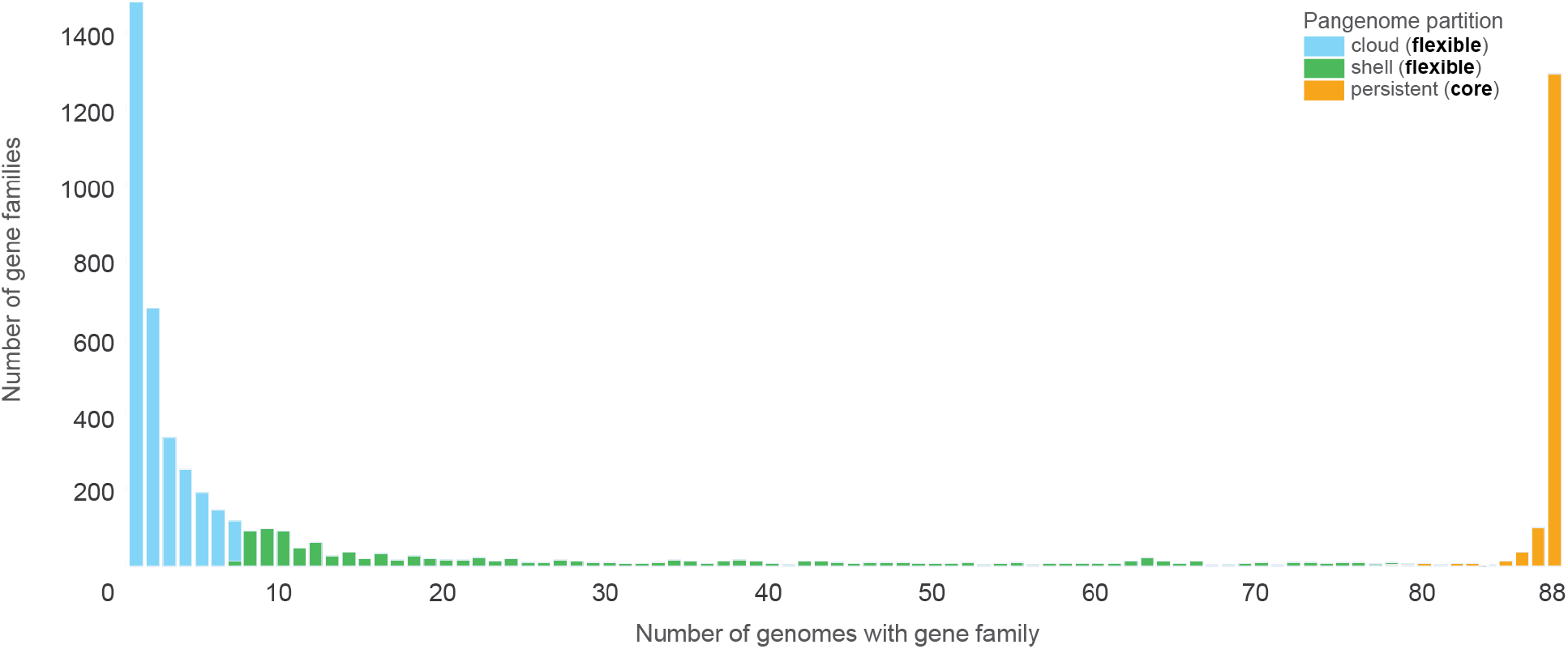
Pangenomic partitioning of all 88 *Pg* strains reveals an abundance of flexible genes. Plot indicates the number of gene families occurring in each of a given number of *Pg* genomes, from gene families that occur in only one genome to gene families that are found in all 88 (as predicted by PPanGGOLiN^53^ using clustered proteins for *Pg*_set88). Light blue and green bars represent counts of gene families with “cloud” and “shell” designations by PPanGGOLiN^53^ (combined and referred to in the text as making up the “flexible” pangenome), respectively, while orange bars represent “persistent” designations (referred to in the text as making up the “core” pangenome).

**Supplementary Table 1.** Summary information describing differences between sets of *Pg* genomes used in this study, with *Pg_set_88* including all genomes and *Pg*_set_79 including only representatives of each strain to eliminate inflation of feature counts in various analyses resulting from inclusion of near-identical genomes.

|  | *Pg*_set_88<br>All genomes in the study | *Pg*_set_79<br>Only representatives of each strain |
| --- | --- | --- |
| # of *Pg* genomes considered | 88 | 79 |
| # genes total | 180,652 | 162,242 |
| # genes included in PPanGGOLiN msmseqs2 protein clusters | 173,833 | 156,170 |
| # of protein clusters | 5,789 | 5,745 |
| # of core protein clusters (“persistent”) | 1,476 | 1,476 |
| # of flexible protein clusters (“shell” + “cloud”) | 4,313 | 4,269 |
| # of shell protein clusters | 1,112 | 1,111 |
| # of cloud protein clusters | 3,201 | 3,158 |
| # of genes in protein clusters in RGPs | 26,659 | 23,938 |
| # of protein clusters in RGPs | 3,097 | 3,060 |
| # of core protein clusters (“persistent”) in RGPs | 84 | 82 |
| # of flexible protein clusters (“shell” + “cloud”) in RGPs | 3,013 | 2,978 |
| # of shell protein clusters in RGPs | 908 | 903 |
| # of cloud protein clusters in RGPs | 2,105 | 2,075 |
| # of flexible protein clusters (“shell” + “cloud”) in RGPs with defense genes and that are not in prophage regions | 1,650 | 1,636 |
| # of protein clusters that are in prophage regions | 353 | 351 |

**Supplementary Figure 6.**
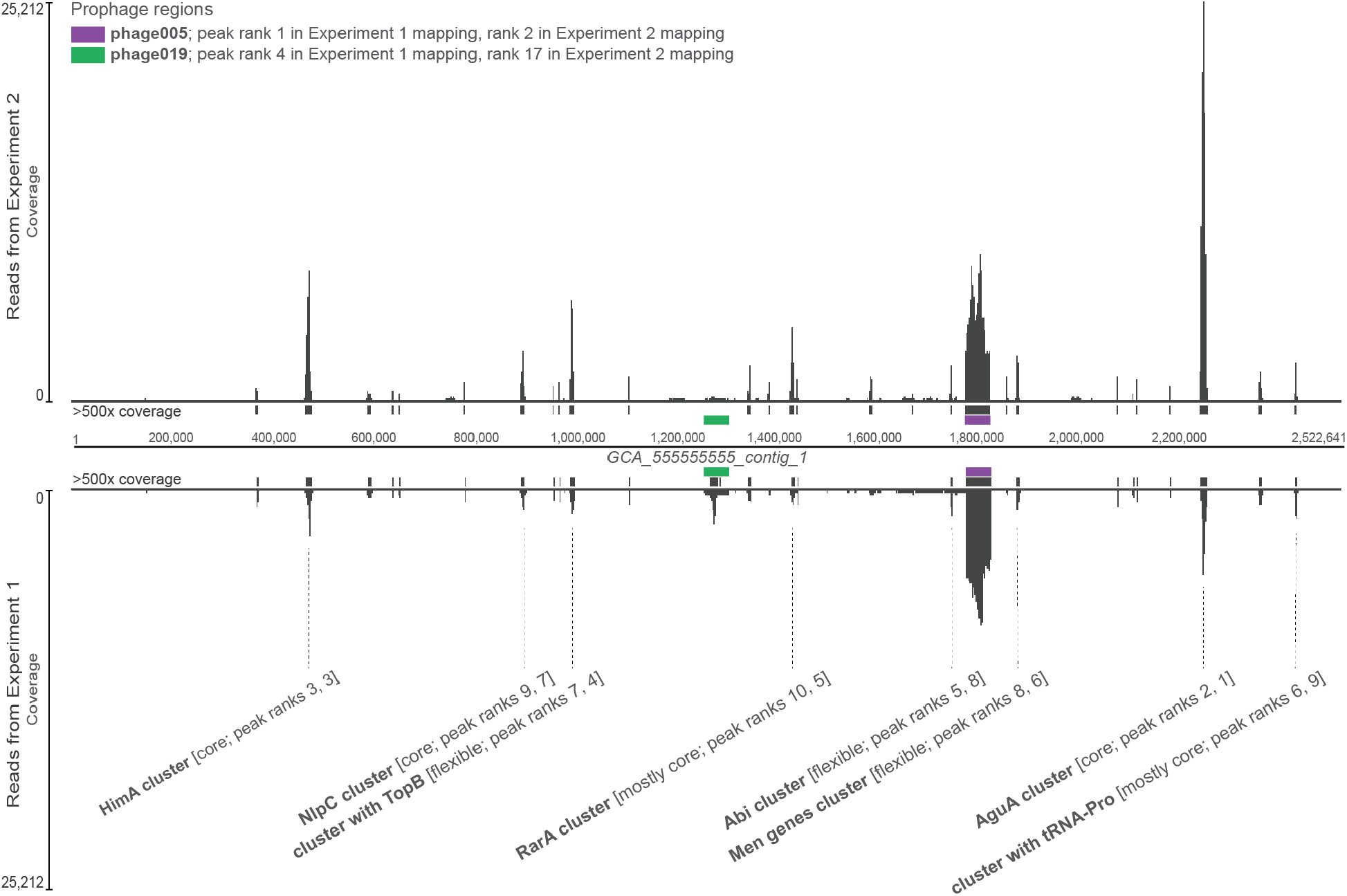
Read mapping of filtered, nuclease-treated supernatants reveals the presence of protected, extracellular DNA in *Pg* cultures. Mapping of Illumina sequencing reads from DNA extracted from cell-free, nuclease-treated, ultracentrifuge-pellets of supernatants of ATCC 49417 *Pg* cultures. Bottom mapping represents reads from “Experiment 1”, which used supernatants from a 19-day old culture; an aliquot of the same culture at a younger age was used to obtain cell pellets from which assembly GCA_444444444 was produced. Top mapping represents reads from “Experiment 2”, which used supernatants from a 20-day old culture; an aliquot of the same culture at a younger age was used to obtain cell pellets from which assembly GCA_555555555 was produced. Both cultures were struck from glycerols originally derived from the same parent glycerol. Reads from both experiments were mapped onto the closed GCA_555555555 assembly and show coverage spikes along the genome (dark gray plots). Regions with greater than 500x coverage are marked by dark gray bars along the length of the reference, regions encoding phage005 and phage019 and marked by purple and green bars, respectively. Select additional peaks of high coverage are also shown, with clusters of elevated coverage named for the gene of highest coverage within the cluster or, where the peak gene is a hypothetical, with the name indicating another gene of known function nearby in the cluster. Mean coverage data for each protein-coding gene in the reference assembly is provided in Supplementary Data 10.

## Materials & Methods

### Bacterial strains and growth conditions

*Porphyromonas gingivalis* isolates Bg4, A7436-C, and HG1691-OLD were shared by Robert E. Schifferle (University at Buffalo, Buffalo, NY) and isolate ATCC 49417 was purchased from the American Type Culture Collection (Manassas, VA). Glycerol stocks of each strain were streaked onto BHI blood agar plates (adapted from Floyd Dewhirst, Forsyth Institute, MA) [Brain Heart Infusion (BD Difco Bacto 237500) - 37 g/L, sodium bicarbonate (JT Baker 3506-01) - 1 g/L, yeast extract (VWR J850) - 5 g/L, and L-cysteine (Sigma-Aldrich C7352) - 0.5 g/L; then supplemented (post-autoclaving) with hemin (Sigma-Aldrich H9039) - 1 mL (5 mg/mL stock concentration), 1,4-dihydroxy-2-naphthoic acid (TCI D2296) - 10 mL (0.1 mg/mL stock concentration), and defibrinated sheep blood (Bio Link Inc) - 53 mL]. After 5 days (6, for ATCC 49417) of anaerobic incubation at 37°C in a GasPak jar with an EZ Anaerobe Container System Sachets with Indicator (BD BBL), multiple colonies from each plate were inoculated into two (three, for ATCC 49417) 100 mL volumes, respectively, of pre-reduced, modified ATCC 2722 broth [Tryptic Soy Broth (Soybean-Casein Digest Medium) (BD Bacto 211825) - 30 g/L, yeast extract (VWR J850) - 5 g/L, and L-cysteine (Sigma-Aldrich C7352) - 0.5 g/L; then supplemented (post-autoclaving) with hemin (Sigma-Aldrich H9039) - 1 mL (5 mg/mL stock concentration) and 1,4-dihydroxy-2-naphthoic acid (TCI D2296) - 10 mL (0.1 mg/mL stock concentration) and anaerobically incubated at 37°C in a Coy chamber (supplied with 5% CO2, 5% H2, 90% N2) or GasPak jar with an anaerobic sachet (for ATCC 49417). As previously noted^83,84^, addition of DHNA was found to be especially beneficial in supporting growth of *Pg* strains.

### Bacterial sequencing and genome assembly

At 2 days post-inoculation of Bg4, A7436-C, and HG1691-OLD cultures (1 day, for ATCC 49417), 1.5 mL from each of 2 replicate culturesOk, per strain, was pooled and centrifuged (Beckman Coulter Allegra X-22R Centrifuge with F2402H rotor) at 5,000 x g (4°C) for 10 min to pellet the cells (with the exception of A7436-C which required an additional 15 min of centrifugation at 7,500 x g). After the centrifugation was complete, the supernatants were removed and the pelleted cells were frozen on dry ice and stored at −80°C. The pellets were extracted and sequenced by the SeqCenter (Pittsburgh, PA) using both Illumina and Nanopore. As reported by SeqCenter: For Illumina sequencing, sample libraries were prepared using the Illumina DNA Prep kit and IDT 10bp UDI indices and sequenced on an Illumina NextSeq 2000 (2×151bp reads). Demultiplexing, quality control and adapter trimming was performed with bcl-convert (v3.9.3). For Nanopore sequencing, runs were on a MinION with an R9 pore type (R9.4.1), base calling was done in high accuracy mode, and Guppy v5.0.16 was used. Genome assemblies were then performed in house from sequences of each culture (Bg4=GCA_111111111.1, A7436-C=GCA_222222222.1, HG1691-OLD=GCA_333333333.1, and ATCC 49417=GCA_555555555.1; all GCAs representing placeholders until assignment of final identifiers by NCBI, see Data Availability statement for link to sequences). In brief, Illumina read quality control was performed by fastp v.0.23.2^85^ (https://github.com/OpenGene/fastp) [fastp --thread $threadsFastp --in1 $internalWorkingIlluminaBax/“$1”\_R1.fastq.gz* --in2 $internalWorkingIlluminaBax/“$1”\_R2.fastq.gz* --out1 “$1”\_R1.fastp.fastq.gz --out2 “$1”\_R2.fastp.fastq.gz --unpaired1 “$1”\_R1.fastp.solo.fastq.gz --unpaired2 “$1”\_R2.fastp.solo.fastq.gz --json “$1”.fastp.json --html “$1”.fastp.html], while Nanopore read quality control was performed by Filtlong v.0.2.1^86^ (https://github.com/rrwick/Filtlong) considering a minimum length threshold of 1000 and keeping 95% of the best reads [filtlong $filtlongMinLength $filtlongKeepPercent “$1”\_nanopore.porechop.noMiddle.fastq.gz > $internalWorkingNanoporeBaxPorechopFiltlong/“$1”\_nanopore.porechop. noMiddle.filtlong.fastq.gz] and Porechop v.0.2.4^87^ (https://github.com/rrwick/Porechop), discarding reads with middle adaptors [porechop -i “$1”\_nanopore.fastq.gz -o ../seq_MiGS_nanopore_bax_porechop/“$1”\_nanopore.porechop.noMiddle.fastq.gz --discard_middle --threads $threadsPorechop]. Hybrid assemblies with the optimized Illumina and Nanopore reads were produced with Unicycler v.0.5.0^88^ (https://github.com/rrwick/Unicycler) under default parameters [unicycler −1 $internalWorkingIlluminaBaxFastpTrims/“$1”*R1.fastp.fastq.gz −2 $internalWorkingIlluminaBaxFastpTrims/“$1”*R2.fastp.fastq.gz −l $internalWorkingNanoporeBaxPorechopFiltlong/$1\_nanopore.porechop.noMiddle.filtlong.fastq. gz -o $unicyclerDIR/$1 -t $threadsUnicycler]. The hybrid assemblies were then polished by Polypolish v.0.5.0^89^ (https://github.com/rrwick/Polypolish) [polypolish assembly.fasta alignments_1.sam alignments_2.sam > polypolish.fasta] with Illumina read alignments by BWA v.0.7.17^90^ (https://github.com/lh3/bwa) [bwa index assembly.fasta bwa mem -t $threadsBwa -a assembly.fasta; $internalWorkingIlluminaBaxFastpT rims/“$1”\_R1.fastp.fastq.gz > alignments_1.sam bwa mem -t $threadsBwa -a assembly.fasta; $internalWorkingIlluminaBaxFastpTrims/“$1”\_R2.fastp.fastq.gz > alignments_2.sam] and MaSuRCA v.4.0.9^91^ (https://github.com/alekseyzimin/masurca) (using POLCA^92^) [polca.sh -a polypolish.fasta -r “$internalWorkingIlluminaBaxFastpTrims/$1*R1.fastp.fastq.gz $internalWorkingIlluminaBaxFastpTrims/$1*R2.fastp.fastq.gz” -t $threadsPolca -m 10G].

### Phage sequencing and read mapping

*For strains ATCC 49417, Bg4, and HG1691-OLD:* At 19 days post-inoculation (20 days, for ATCC 49417), 184 mL (231 mL, for ATCC 49417) from each replicate culture (same as those previously described for bacterial sequencing) were pooled, per strain, and filtered using a 0.22 μm filter system (Corning) to remove the cells. Phages were ultracentrifuged (Beckman Coulter Optima XE-90 Ultracentrifuge with SW 32 Ti rotor) at 174,900 x g (22°C) for 1 hr (repeated until each culture was completely pelleted by removing the supernatant and refilling the tubes, followed by a rinse centrifugation with SM buffer) in Ultra-Clear centrifuge tubes (Beckman) pre-rinsed with sterile distilled water. After the centrifugation, the supernatant was removed and the pellet was allowed to resuspend overnight (4 days, for ATCC 49417) at 4°C in enough SM buffer to cover the pellet. The next morning, the pellets were rocked for approximately 2 hr at room temperature (~22°C), the resuspended pellet was removed, and each tube washed with approximately 450 μL SM buffer for a total volume of ~900 μL for each strain to be used in the phage DNA extraction protocol modified from Jakočiūnė and Moodley 2018^93^. In brief, the resuspended pellets were each split into two 450 μL samples. The unprotected nucleic acids were removed by adding 50 μL of 10x Turbo DNase Buffer (Qiagen), 5 μL of 2U/μL Turbo DNase (Qiagen), and 1 μL of 10 mg/mL RNase A (Qiagen), then incubating at 37°C for 1.5 hr without shaking. The nucleases were then denatured by simultaneously adding 20 μl of 0.5 M EDTA and 57 μL of 20 mg/μL Proteinase K (Qiagen) and incubating at 56°C for 2 hr, vortexing every 20 min (an additional 57 μL of Proteinase K was added at 100 (60, for ATCC 49417) min because the sample was still cloudy). The once-protected DNA was then extracted using the DNeasy Blood & Tissue Kit (Qiagen). First, an equal volume of AL Buffer (Qiagen) was added to each sample, these were then vortexed and incubated at 70°C for 10 min. After incubation, the same volume of 100% ethanol was added and the samples were vortexed. The samples were transferred into DNeasy Mini spin columns (Qiagen) and centrifuged (Beckman Coulter Allegra X-22R Centrifuge with F2402H rotor) at 6,000 x g (22°C) for 1 min. This was repeated several times until all of each sample was run through the spin column. Next, 500 mL of Buffer AW1 (Qiagen) was added to the spin columns which were then centrifuged at 6,000 x g (22°C) for 1 min. Then, 500 mL of Buffer AW2 (Qiagen) was added to the spin columns which were then centrifuged at 20,000 x gf (22°C) for 3 min. Lastly, 40 mL of AE Buffer (Qiagen) was added directly onto the spin column membrane and let incubate at room temperature for 1 min prior to centrifugation at 6,000 x g (22°C) for 1 min. The collected DNA was then shipped to SeqCenter (Pittsburgh, PA) for Illumina sequencing. The sequenced reads were then mapped back to the bacterial assembly produced from their respective culture with BWA v.0.7.17^90^ [bwa mem $base *R1_001.fastq.gz *R2_001.fastq.gz | samblaster -e -d $base.disc.sam -s $base.split.sam | samtools view -Sb - > $base.out.bam 2>&1 | tee $base.samblaster.output.log] and using SAMBLASTER v.0.1.26^94^ (https://github.com/GregoryFaust/samblaster). To determine the average coverage per gene, the mapped reads were aligned to a bed file of protein-coding regions with bedtools v.2.30.0^95^ (https://github.com/arq5x/bedtools2) [bedtools coverage -a GCA_555555555.MEGA.bed -b GCA_555555555.out.bam -mean > GCA_555555555.bedtools.out]. *For strain A7436-C:* A7436-C cell-free, nuclease-protected DNA was extracted, sequenced, and mapped similarly to that previously described, with the few minor exceptions listed here. First, three-100 mL pre-reduced, modified ATCC 2722 broths were inoculated with multiple colonies from a 5-day old streak on a BHI blood agar plate. After 17 days of anaerobic incubation in a Coy chamber, the cultures were pooled and left to filter by gravity for 7 days. After the filtration, the pellet was eluted for 4 days at 4°C. Secondly, during the nuclease deactivation, no additional Proteinase K was added to the incubation which lasted 1.5 hr (due to the sample being clear).

### Additional bacterial and phage sequencing

To highlight reproducibility we note that an additional ATCC 49417 bacterial culture (derived from the same parent glycerol stock that produced ATCC 49417 assembly GCA_555555555.1) was sequenced and assembled (GCA_444444444.1). Whereas the GCA_555555555.1 assembly yielded a single closed contig, the GCA_444444444.1 yielded an assembly where the prophage region was represented as an independent contig. This difference is interpreted as reflecting differential relative abundances of extra-chromosomal and integrated versions of one of the prophages in the two cultures. Filtered supernatants of the additional ATCC 49417 cultures were also ultracentrifuged, nuclease treated, extracted, Illumina sequenced, and the reads mapped onto the GCA_555555555.1 assembly, and found to have similar profiles of nuclease-protected DNA between cultures (Supplementary Fig. 6).

### Electron microscopy of active phages

To determine if active phages are produced from ATCC 49417 lysogens, two different samples were imaged via transmission electron microscopy. The first sample was material from the cell-free ATCC 49417 ultracentrifuged pellet (same as which was described for phage sequencing), prior to nuclease treatment, that is shown in Fig. 6A, B. The second sample was a separate subculture of ATCC 49417 (from the same stock that also gave rise to the subculture used in the bacterial and phage sequencing), that is shown in Fig. 6C, D. This sample was struck out from a glycerol stock onto a BHI blood agar plate supplemented with 100 μL of a 10:1 dHNA-hemin stock mix and incubated anaerobically at 37°C. After 5 days of incubation, three 10 μL inoculation loops through the tail end of the streak were inoculated into 200 mL of modified ATCC 2722 broth and was incubated anaerobically for 3 days. Both samples were identically prepared on formvar/carbon film 200 mesh copper grids (Ted Pella 01803-F). First, the grids were glow discharged for 5 seconds to improve their hydrophilicity prior to adding 5 μL of the sample. After 30 seconds, the sample was drawn off and the grids were rinsed with 5 μL of nuclease-free water (Invitrogen AM9938). The water was then drawn off and the rinse was repeated. Lastly, after the water from the second rinse was drawn off, 5 μL of 1% uranyl acetate in water (Electron Microscopy Sciences 22400-1) was added to the grid, then immediately drawn off to let the grid air dry for 20 minutes. The grids were imaged at University at Buffalo’s Electron Microscopy Core Lab (Jacobs School of Medicine and Biomedical Sciences, Buffalo, NY) on a Hitachi HT7800 high resolution 120kV transmission electron microscope with a Gatan Rio 16 CMOS camera capturing 4k x 4k pixel images.

### Selection and curation of *Pg* genomes used in bioinformatic analyses

We sought to obtain a comprehensive set of high quality *Pg* genomes, and ultimately defined two sets for analyses in this work: *Pg*_set88 and *Pg*_set79. We considered three sources of Pg genome assemblies for inclusion in this study, as follows. First, we included four of the five genomes sequenced and assembled in house, as described above, excluding GCA_444444444.1 from *Pg*_set88 as a duplicate assembly of ATCC 49417. Second, we considered *Pg* assemblies available in NCBI GenBank (88 assemblies initially). To ensure that our collection of GenBank-derived assemblies was comprehensive and free of mislabeled strains (false *Pgs*) we obtained all assemblies for the genus *Porphyromonas* from GenBank and generated whole genome phylogenies using BacSort (https://github.com/rrwick/Bacsort) with the combined FastANI^96^ and Mash^97^ approach to generate a distance matrix and tree for visualization with phyloXML^98^ and Archaeopteryx (https://www.phylosoft.org/archaeopteryx/). We found that all strains labeled in GenBank as *Pg* were members of a single monophyletic clade containing no unlabeled or mislabeled strains, with *P. gulae* the nearest neighboring clade. Four metagenome-derived assemblies were excluded from *Pg*_set88 on the basis of each of their total sizes being <2Mb. Finally, we considered *Pg* assemblies available in GenBank and re-assembled in house (as described above) to explore potential for improved assemblies facilitating detection of phages otherwise split across multiple contigs. In exploratory analyses we found that re-assemblies did not recover additional prophage regions and thus these were also excluded. Thus *Pg*_set88 included four genomes sequenced in house and 84 genomes from GenBank. To reduce inflation of feature counts in various analyses resulting from inclusion of near-identical genomes, we assign one assembly as the “primary” assembly in all cases where we have multiple assemblies with the same strain name (A7A1_28, ATCC 33277, ATCC 49417, TDC60, W50, W83), or which are known laboratory-derivatives (e.g. W50/BE1, W50/BR1, A7436C). The set of primary assemblies is identified as *Pg*_set79. Primary assemblies were selected as those with the smaller number contigs, and if the number of contigs was the same then the more recent assembly was selected. In the case of genomes representing derivatives, the parent strain was assigned as the primary assembly. Information on all sequences considered is available in Supplementary Data 1.

### Reference phylogeny, gene annotation, and pangenome analysis of *Pg* genomes

To obtain a reference phylogeny for use throughout our study we used RiboTree (https://github.com/philarevalo/RiboTree), which considers single copy ribosomal proteins^99^ with default parameters, using *P. gulae* assembly (GCA_000768765.1) as an outgroup. To standardize formatting and functional annotation across all *Pg*_set88 assemblies, we used Bakta^54^ (https://github.com/oschwengers/bakta) with default parameters [bakta --db $baktaDbDIR --verbose --output $baktaOutDIR/$base --prefix $base --min-contig-length 200 --genus $genus --species $species --gram $baktaGram --compliant --threads $threadsBakta --keep-contig-headers $baxAssembliesTask020DIR/$1]. To define pangenome partitions and regions of genome plasticity (RGPs, runs of predominantly flexible genes) we used PPangGGOLiN^53^ (https://github.com/labgem/PPanGGOLiN) with Bakta^54^ gene calls and otherwise default parameters [ppanggolin all --anno ORGANISMS_FASTA_LIST --cpu $threadsPpanggolin].

### Identification of CRISPR-Cas and other defense systems in *Pg* genomes

To identify putative CRISPR-Cas systems, *Pg*_set88 genomes were evaluated using CRISPRCasTyper^36^ command line CCTyper v.1.6.4 [cctyper $baxBaktaGenomesStdsDIR/$1 $cctyperOutDIR/$base --keep_tmp] and webserver https://crisprcastyper.crispr.dk) with default parameters. Full summary data are available in Supplementary Data 4. CCTyper^36^ identifies *cas* operons (certain and putative) and CRISPR arrays, annotates each on the basis of repeat and *cas* gene similarity to known systems, and combines this information to identify high confidence CRISPR-Cas systems. Our summary data regarding the number of high confidence CRISPR-Cas systems in the 88 *Pg* genomes excludes cases where the *cas* operon classification was ambiguous and where *cas* genes or CRISPR arrays were identified but could not be readily linked to one another, in some cases likely due to fragmented assemblies. Our analysis of the total number of unique spacers, and the proportion that could be mapped to phages, includes data from all identified CRISPR arrays, including those to which *cas* operons could not be linked, and was performed as follows. All spacers were identified to classes on the basis of the CRISPR-Cas operon assignment by command line CCTyper^36^, or by the subsequent classification of the consensus repeat for the array by the CCTyper^36^ webserver (accessed 11/13/2022), which offers a more frequently updated repeat classification model. Final standardized sequence orientations of repeats and spacers in all arrays were determined on the basis of the strand of the associated *cas* operon interference module or based on identical (direct or by reverse complement) consensus repeats in systems with assigned directions. In cases of Type I-B, I-C, III-B, and VI-B2 systems the directionality of the array repeats and spacers was set the same as for the *cas* operon, whereas for Type II-C systems the directionality was set to be the reverse^100^. Using this approach yielded a total of 4,016 unique spacer sequences (including those with Ns), 4,015 when considering reverse complements. To identify candidate non-CRISPR-Cas defense systems we also annotated all *Pg*_set88 genomes using the PADLOC^55^ webserver (https://padloc.otago.ac.nz/padloc/) with PADLOC-DB v1.4.

### Mapping of CRISPR spacer hits to bacterial and phage genomes

To map CCTyper^36^ spacers to bacterial genomes and extracted phages (see below) we use Bowtie v.1.1.1^76^, considering separately the number of hits for both 0 and 1 mismatch to the target sequence (as shown in Fig. 3) [0 mismatch: bowtie -a -v 0 $base -f $cctyperStdOspacers/$cctyperSpacersStdONAME --sam > $base.cctyper.sam; 1 mismatch: bowtie -a -v 1 $base -f $cctyperStdOspacers/$cctyperSpacersStdONAME --sam > $base.cctyper.sam]. To also allow evaluation of hits to non-phage sequences, we additionally mapped coverage on a per gene basis in bacterial genomes using the BEDTools^95^ annotate function with default settings. Exploratory analyses of differences in strand level mapping of spacers from Class 1 systems [where crRNAs bind DNA (Types I-B, I-C, and III-B)], versus Class 2 systems [where crRNAs bind mRNA (Types II-C and VI-B2)]^42^,^101^, showed no consistent patterns. Both direct and reverse complement mappings were therefore counted for all spacers. We note that even where systems are known to have strand preferences there is generally also representation of the other strand among targets in the spacer array^102^, perhaps as a result of trans interactions between different systems^44^. To identify potential matches to phages with conserved protein sequences but divergent nucleotide sequences translated spacer sequences were mapped to phage proteins using SpacePHARER^37^ (https://github.com/soedinglab/spacepharer) with default parameters [spacepharer predictmatch $spacepharerSpacerDbDIR/spacerSetDB $spacepharerPhageDbDIR/phageSetDB $spacepharerPhageDbDIR/controlPhageSetDB $spacepharerOutDIR/2022.320_spacepharerOutput.tsv $spacepharerOutDIR/tmpFolder]. To determine whether the *Pg* phages potentially have alternate hosts we used CRISPROpenDb^41^ (https://github.com/edzuf/CrisprOpenDB) to map spacer sequences harvested from other species of bacteria to all the bacterial genomes [python CL_Interface.py -i $baxBaktaGenomesStdsSplitsDIR/$1 --mismatch 0 --num_threads $threadsCrisprOpenDB --report > $baxCrisprOpenOutDIR/“$base”.spacerHits], as well as the extracted phages separately [python CL_Interface.py -i $importPhageFastas/$1 --mismatch 0 --num_threads $threadsCrisprOpenDB --report > $importPhageCOdbDIR/$base/$base.spacerHits].

### Quantification of prophage and defense system contributions to the *Pg* pangenome

The contribution of prophage genes to the *Pg* pangenome was determined by identifying all bacterial gene families (with prefix mmseq.000837.22272) occurring in prophage regions. The proportion of the pangenome associated with defense was determined by identifying all regions of genome plasticity (RGPs, as defined by PPanGGOLiN^53^, and representing runs of predominantly flexible genes) that contained any of the following: any gene families for which any member was identified by CCTyper^36^ as part of a CRISPR-Cas systems, any gene families for which any member was identified by PADLOC^55^ as a defense system, any gene families not captured by the aforementioned tools but for which any member was annotated with functions containing defense-function related keywords (e.g. cas, CRISPR, restriction, abortive infection, Abi, death on curing, addiction, toxin/antitoxin). One gene family identified as defense related but annotated as a transposon was excluded (mmseq.000837.22272.2484). Any RGPs containing any predicted defense-related proteins were considered as potential defense islands or elements, and all non-core gene families in all of these RGPs were counted toward the estimate of total gene families represented by defense islands or elements.

Identification of prophages in Pg genomes. To identify prophages in Pg_set88 genomes we combined multiple complementary approaches and used Geneious Prime versions 2023.0.1 and 2022.2.2 (Biomatters Ltd.) to view all results together and manually curate prophage boundaries. As initial exploratory investigations revealed that some prophage regions were fragmented, our analysis of the Pg genomes included a set of “fusion contigs” generated by manual targeted curation to identify contigs encoding genes for which there was evidence of terminal overlap. Fusion contigs were generated for 3 strains (as noted in Supplementary Data 1), with these contigs renamed with terminal “9”s to indicate their having been updated from their original assemblies (e.g. JAEMBP01999999.1 fusion contig from contigs JAEMBP010000058.1 and JAEMBP010000009.1). All Pg_set88 genomes, including updated fusion contigs, were then searched for predicted prophage regions using CenoteTaker2^79^ (https://github.com/mtisza1/Cenote-Taker2), VIBRANT^81^ (https://github.com/AnantharamanLab/VIBRANT), PhageBoost^80^ (https://github.com/ku-cbd/PhageBoost), VirSorter2^77^ (https://github.com/jiarong/VirSorter2) post-processed with CheckV^78^ (https://bitbucket.org/berkeleylab/CheckV/src), and Inovirus detector^82^ (https://github.com/simroux/Inovirus). To facilitate determination of nucleotide-level boundaries of phage regions we used an all-by-all BLAST of all Pg_set88 genomes [blastn -task megablast -query $baxBaktaGenomesStdsDIR/$1 -parse_deflines -db $intrapopBlastDbDIR/baxBaktaStdGenomes.all -perc_identity $pid -outfmt “6 qseqid pident length qstart qend sseqid” -word_size $word -num_threads $threadsBlast > $base\_v_all.tsv]. As described above, to facilitate detection of regions targeted by CRISPR spacers we identified all CRISPR array spacers in Pg_set88 and mapped these back to all Pg_set88 genomes using Bowtie^76^, and we also identified all sites targeted by CRISPR spacers encoded in other bacterial species using CRISPROpenDB^41^. All contigs with predicted prophage regions were then imported into Geneious and evaluated together with tracks showing: pangenome partition information for all bacterial genes, all-by-all BLAST results, and CCTyper^36^ and CRISPROpenDB^41^ spacer mappings. Repeats surrounding candidate regions were next identified using the Geneious Repeat Finder v1.0.1, and final boundaries defined based on identification of bounding repeats proximal to conserved BLAST hit edges (identifying regions commonly showing gaps in Pg genomes) and corresponding to regions identified as flexible pangenome partitions. This approach readily revealed boundaries for tRNA-inserting phages, which generally had bounding repeats of ≥13bp (with one repeat being part of the phage genome), however for the transposable phages additional curation was needed and included extraction and alignment of candidate regions to identify conserved termini and short 4bp bounding repeats (both outside the boundaries of the phage genome).

### Prediction of *Pg* phage genes

Exploratory analyses revealed that predicted open reading frames in prophage regions were inconsistently identified both by Bakta^54^ in the original bacterial genome annotations, and by Prodigal^103^ run separately on only the extracted prophage regions. Therefore, all prophage regions were re-analyzed with CenoteTaker2^79^, which provides excellent functional annotation of open reading frames predicted using PHANOTATE^104^, a gene caller optimized for phage genes [python $cenotePATH -c $importPhageFastas/$1 -r $run -m 225 -t $threadsCenote -hh hhsearch -p false -am True --wrap False -db standard 2>&1 | tee $run.cnt2.output.log]. All protein coding genes predicted PHANOTATE^104^ were clustered using mmseqs2^105^ [mmseqs easy-cluster cenote.geneious.ALL.prots.fasta clusterResult mmseqsTMP --createdb-mode 0 --min-seq-id 0.8 -c 0.8 --cov-mode 1] and thus phage regions have two sets of protein clusters in our study, those derived from the original Bakta^54^ gene calls in the bacterial genomes (identified with the prefix mmseq.000837.22272), and those derived from PHANOTATE^104^ gene calls on the extracted phage genomes (identified with the prefix mmseq.010239.22272).

### Annotation of *Pg* phage genes

All phage gene annotations were performed on PHANOTATE^104^ derived gene calls as described above. Phage proteins were annotated for predicted function by comparison to the PHROGS^106^ v4 database (https://phrogs.lmge.uca.fr/index.php) using 3 iterations of HHblits^107^ [hhblits -i $file -d $phrogsHHsuiteDIR/phrogs -n 3 -o $file\_VS_phrogs.out -blasttab $file.HHBLITS.tsv_file -cpu $threadsHHblits] and allowing automatic assignment to top hit annotations and categories with a bitscore of >30, where not superceded by another annotation. Additional annotations included those provided by CenoteTaker2^79^, Bakta^54^ using PHANOTATE^104^ gene calls as input, eggNOG-mapper^108,109^ (http://eggnog-mapper.embl.de/), Batch CD-Search^110–114^ (https://www.ncbi.nlm.nih.gov/Structure/bwrpsb/bwrpsb.cgi), Phyre2^115^ (http://www.sbg.bio.ic.ac.uk/phyre2), HHpred through the MPI Bioinformatics Toolkit^116^, SignalP6.0^117^ (https://services.healthtech.dtu.dk/service.php?SignalP-6.0, using “Fast” model option for initial run and “Slow” model option for refining cleavage sites of candidates identified in initial run), and jackhmmer^118^ (https://www.ebi.ac.uk/Tools/hmmer/search/jackhmmer). Candidate anti-CRISPR (acr) genes were predicted using two approaches. First, direct annotation of phage protein coding genes on the PaCRISPR^39^ webserver (https://pacrispr.erc.monash.edu/index.jsp). Second, proteins in the DeepAcr database were mapped to PHROG gene families [prepare database: mmseqs createdb Acr_predictions.fasta Acr_predictions; search database: mmseqs search $phrogsMmseqsDIR/phrogs_profile_db Acr_predictions out_deepacrVphrogs_mmseqs./tmp -s 7], and any *Pg* phage gene that was identified as also mapping to the same PHROG was annotated as an acr. Only phage genes identified through both approaches were ultimately annotated as candidate acrs and colored accordingly in the Fig. 2 phage genome diagrams. Select candidate spanins were identified using tools available on the Center for Phage Technology Galaxy Server^119^ (https://cpt.tamu.edu/galaxy-pub) (run errors resulted in lack of even annotation across all phage genes) and often showed frameshifts from open reading frames identified by PHANOTATE^104^; in addition, information on lipoprotein signal peptides and proximity to other lysis cassette genes such as the endolysin and holin were also considered. Phage morphotypes and head-neck-tail components were predicted using VIRFAM^120^ (http://biodev.cea.fr/virfam/). Except in the case of annotation of anti-crispr proteins, in cases where only a single representative of a protein cluster was annotated (e.g. with Phyre2^115^), annotations from any member were propagated to all other members of the protein cluster and annotations overall were harmonized within protein clusters. All annotations of phage protein coding genes are available in Supplementary Data 3.

### Analysis and visualization of phage genome relatedness

To determine whether any of the phages identified in this work were related to known phages, all were compared to references using ViPTree^121^ (https://www.genome.jp/viptree/). All *Pg* phage sequences, along with nearest neighors on the ViPTree (NC_019490.1, *Riemerella* phage RAP44; NC_047910.1, *Faecalibacterium* phage FP_Epona; NC_062754.1, *Winogradskyella* phage Peternella_1) were then compared using VIRIDIC^122^ (https://rhea.icbm.uni-oldenburg.de/VIRIDIC/) with default parameters [blastn -word_size 7 -reward 2 -penalty −3 -gapopen 5 -gapextend 2; species threshold 95% sequence identity; genus threshhold 70% sequence identity]. Final trees for visualization of phage relationships were generated using VICTOR^74^ using whole genome BLAST distance with concatenated proteins and scaled by the VICTOR^74^ d6 formula, resulting midpoint-rooted trees were visualized with iTOL^123^ (https://itol.embl.de/). Phage genome diagrams were visualized with Clinker^75^ (https://github.com/gamcil/clinker). All final figures assembled using Adobe Illustrator.

### Exploratory mapping of healthy and periodontal disease metagenomes to Pg phages

Illumina reads from publicly available metagenomic samples from a study^124^ of six healthy and seven periodontitis patients were downloaded from the Human Oral Microbiome Database^125^ (https://homd.org/ftp/publication_data/20130522/). The reads from each patient were mapped to each of the 35 *Pg* reference phage genomes using Geneious Prime v.2022.2.2 (Biomatters Ltd.) with the Geneious mapper at default settings, with the exception of mapping multiple best matches to all locations (such that reads mapping to multiple phages would be represented in coverage mappings from each).

### Bioinformatic analyses

Unless otherwise specified above, bioinformatic analyses were conducted on the UB High Performance Compute Cluster using Miniconda (https://docs.conda.io/en/latest/miniconda.html), conda environments (https://docs.conda.io/en/latest/) installed from the Anaconda Package Repository (https://anaconda.org/anaconda/repo), and in house Unix shell script wrappers.

## Supplementary data files

**Supplementary Data 1.** Overview of bacterial and phage genome information

**Supplementary Data 2.** Virulence

**Supplementary Data 3.** Phage protein coding gene annotations

**Supplementary Data 4.** CRISPR-Cas system annotations

**Supplementary Data 5.** CCtyper CRISPR spacer hits to phages

**Supplementary Data 6.** SpacePHARER CRISPR spacer hits to phages

**Supplementary Data 7.** CRISPROpenDB CRISPR spacer hits to phages

**Supplementary Data 8.** Bacterial protein coding gene annotations.

**Supplementary Data 9**. Transposases

**Supplementary Data 10.** Mapping of cell-free nuclease-protected DNA to *Pg* ATCC 49417

## Data availability

All *Pg* genomes sequenced for this work have been deposited to NCBI GenBank BioProject PRJNA874424 (https://www.ncbi.nlm.nih.gov/bioproject/PRJNA874424) with BioSample accession numbers SAMN30559729-SAMN30559733. All other data not already included in Supplementary Data files or described here, as well as any wrapper shell scripts used to run described publicly available bioinformatic tools, are available upon request from the authors. We note that for the initial submission we have used placeholder GCA identifiers (GCA_111111111, GCA_222222222, GCA_333333333, GCA_444444444, GCA_555555555) for new sequences throughout the manuscript, figures, and supplementary files (see Supplementary Data 1), these will be updated throughout once final accession numbers are received, and all sequence files are available at: https://zenodo.org/record/7489347.

## Acknowledgements

We would like to gratefully acknowledge the UB Center for Computational Research for access to the high performance compute cluster and assistance with troubleshooting, and each of the following for helpful discussions, as well as other support as noted: Frank Scannapieco (also for sharing laboratory resources), Robert Schifferle (also for sharing strains), Susan Yost (also for sharing strains), Patricia Diaz, Peter Bush, Matthew Smardz, and Carol Parker.

## Author contributions

Conceptualization, CM, KK; Methodology, CM, EH, DM, KK; Formal Analysis, CM, KK; Investigation, CM, KK; Resources, EH, KK; Data Curation, CM, KK; Writing – Original Draft Preparation, CM, KK; Writing – Review & Editing, CM, EH, FD, JMW, FMS, DM, KK; Visualization, CM, KK; Supervision, Project Administration, and Funding Acquisition, KK.

## References

1. Dewhirst, F. E., Chen, T., Izard, J. & Paster, B. J. The human oral microbiome. Journal of (2010).

2. Hajishengallis, G., Darveau, R. P. & Curtis, M. A. The keystone-pathogen hypothesis. Nat.Rev. Microbiol. 10, 717–725 (2012).

3. Hoare, A. et al. A cross-species interaction with a symbiotic commensal enables cell-density-dependent growth and in vivo virulence of an oral pathogen. ISME J. 15, 1490–1504 (2021).

4. Bosshardt, D. D. & Lang, N. P. The junctional epithelium: from health to disease. J. Dent.Res. 84, 9–20 (2005).

5. Bertozzi Silva, J., Storms, Z. & Sauvageau, D. Host receptors for bacteriophage adsorption. FEMS Microbiol. Lett. 363, (2016).

6. Edlund, A., Santiago-Rodriguez, T. M., Boehm, T. K. & Pride, D. T. Bacteriophage and their potential roles in the human oral cavity. J. Oral Microbiol. 7, 27423 (2015).

7. Willner, D. et al. Metagenomic detection of phage-encoded platelet-binding factors in the human oral cavity. Proc. Natl. Acad. Sci. U. S. A. 108 Suppl 1, 4547–4553 (2011).

8. Jahn, M. T. et al. A Phage Protein Aids Bacterial Symbionts in Eukaryote Immune Evasion. Cell Host Microbe 26, 542–550.e5 (2019).

9. Ly, M. et al. Altered oral viral ecology in association with periodontal disease. MBio 5, e01133–14 (2014).

10. Tylenda, C. A., Kolenbrander, P. E. & Delisle, A. L. Use of bacteriophage-resistant mutants to study Actinomyces viscosus cell surface receptors. J. Dent. Res. 62, 1179–1181 (1983).

11. Tylenda, C. A., Enriquez, E., Kolenbrander, P. E. & Delisle, A. L. Simultaneous loss of bacteriophage receptor and coaggregation mediator activities in Actinomyces viscosus MG-1. Infect. Immun. 48, 228–233 (1985).

12. Delisle, A. L., Donkersloot, J. A., Kolenbrander, P. E. & Tylenda, C. A. Use of lytic bacteriophage for Actinomyces viscosus T14V as a probe for cell surface components mediating intergeneric coaggregation. Infect. Immun. 56, 54–59 (1988).

13. Kolenbrander, P. E. et al. Bacterial interactions and successions during plaque development. Periodontol. 2000 42, 47–79 (2006).

14. Szafrański, S. P., Slots, J. & Stiesch, M. The human oral phageome. Periodontol. 2000 86, 79–96 (2021).

15. Zambon, J. J., Reynolds, H. S. & Slots, J. Black-pigmented Bacteroides spp. in the human oral cavity. Infect. Immun. 32, 198–203 (1981).

16. Sandmeier, H., Bär, K. & Meyer, J. Search for bacteriophages of black-pigmented gram-negative anaerobes from dental plaque. FEMS Immunol. Med. Microbiol. 6, 193–194 (1993).

17. Chen, T., Siddiqui, H. & Olsen, I. In silico Comparison of 19 Porphyromonas gingivalis Strains in Genomics, Phylogenetics, Phylogenomics and Functional Genomics. Front. Cell.Infect. Microbiol. 7, 28 (2017).

18. Haigh, R. D. et al. Draft Whole-Genome Sequences of Periodontal Pathobionts Porphyromonas gingivalis, Prevotella intermedia, and Tannerella forsythia Contain Phase-Variable Restriction-Modification Systems. Genome Announc. 5, (2017).

19. Watanabe, T., Shibasaki, M., Maruyama, F., Sekizaki, T. & Nakagawa, I. Investigation of potential targets of Porphyromonas CRISPRs among the genomes of Porphyromonas species. PLoS One 12, e0183752 (2017).

20. Watanabe, T. et al. CRISPR regulation of intraspecies diversification by limiting IS transposition and intercellular recombination. Genome Biol. Evol. 5, 1099–1114 (2013).

21. Solbiati, J., Duran-Pinedo, A., Godoy Rocha, F., Gibson, F. C., 3rd & Frias-Lopez, J. Virulence of the Pathogen Porphyromonas gingivalis Is Controlled by the CRISPR-Cas Protein Cas3. mSystems 5, (2020).

22. Yost, S., Duran-Pinedo, A. E., Teles, R., Krishnan, K. & Frias-Lopez, J. Functional signatures of oral dysbiosis during periodontitis progression revealed by microbial metatranscriptome analysis. Genome Med. 7, 27 (2015).

23. Pride, D. T. et al. Evidence of a robust resident bacteriophage population revealed through analysis of the human salivary virome. ISME J. 6, 915–926 (2012).

24. Kuzio, J. & Kropinski, A. M. O-antigen conversion in Pseudomonas aeruginosa PAO1 by bacteriophage D3. J. Bacteriol. 155, 203–212 (1983).

25. Tsao Yu-Fan et al. Phage Morons Play an Important Role in Pseudomonas aeruginosa Phenotypes. J. Bacteriol. 200, e00189–18 (2018).

26. Sandulache, R., Prehm, P. & Kamp, D. Cell wall receptor for bacteriophage Mu G(+). J.Bacteriol. 160, 299–303 (1984).

27. Bochtler, M. et al. The Bacteroidetes Q-Rule: Pyroglutamate in Signal Peptidase I Substrates. Front. Microbiol. 9, 230 (2018).

28. Song, S. & Wood, T. K. A Primary Physiological Role of Toxin/Antitoxin Systems Is Phage Inhibition. Front. Microbiol. 11, 1895 (2020).

29. Srikant, S., Guegler, C. K. & Laub, M. T. The evolution of a counter-defense mechanism in a virus constrains its host range. Elife 11, (2022).

30. Guo, Y. et al. RalR (a DNase) and RalA (a small RNA) form a type I toxin-antitoxin system in Escherichia coli. Nucleic Acids Res. 42, 6448–6462 (2014).

31. Rousset, F. et al. Phages and their satellites encode hotspots of antiviral systems. Cell Host Microbe 30, 740–753.e5 (2022).

32. Jørgensen, M. G., Pandey, D. P., Jaskolska, M. & Gerdes, K. HicA of Escherichia coli defines a novel family of translation-independent mRNA interferases in bacteria and archaea. J. Bacteriol. 191, 1191–1199 (2009).

33. Li, G. et al. Identification and Characterization of the HicAB Toxin-Antitoxin System in the Opportunistic Pathogen Pseudomonas aeruginosa. Toxins 8, 113 (2016).

34. Kurata, T. et al. A hyperpromiscuous antitoxin protein domain for the neutralization of diverse toxin domains. Proc. Natl. Acad. Sci. U. S. A. 119, (2022).

35. Chen, T. & Olsen, I. Porphyromonas gingivalis and its CRISPR-Cas system. J. Oral Microbiol. 11, 1638196 (2019).

36. Russel, J., Pinilla-Redondo, R., Mayo-Muñoz, D., Shah, S. A. & Sørensen, S. J. CRISPRCasTyper: Automated Identification, Annotation, and Classification of CRISPR-Cas Loci. CRISPR J 3, 462–469 (2020).

37. Zhang, R. et al. SpacePHARER: Sensitive identification of phages from CRISPR spacers in prokaryotic hosts. Bioinformatics (2021) doi:10.1093/bioinformatics/btab222.

38. Watters, K. E., Fellmann, C., Bai, H. B., Ren, S. M. & Doudna, J. A. Systematic discovery of natural CRISPR-Cas12a inhibitors. Science 362, 236–239 (2018).

39. Wang, J. et al. PaCRISPR: a server for predicting and visualizing anti-CRISPR proteins. Nucleic Acids Res. 48, W348–W357 (2020).

40. Wandera, K. G. et al. Anti-CRISPR prediction using deep learning reveals an inhibitor of Cas13b nucleases. Mol. Cell 82, 2714–2726.e4 (2022).

41. Dion, M. B. et al. Streamlining CRISPR spacer-based bacterial host predictions to decipher the viral dark matter. Nucleic Acids Res. 49, 3127–3138 (2021).

42. Adler, B. A. et al. Broad-spectrum CRISPR-Cas13a enables efficient phage genome editing. Nat Microbiol (2022) doi:10.1038/s41564-022-01258-x.

43. VanderWal, A. R., Park, J.-U., Polevoda, B., Kellogg, E. H. & O’Connell, M. R. CRISPR-Csx28 forms a Cas13b-activated membrane pore required for robust CRISPR-Cas adaptive immunity. bioRxiv 2021.11.02.466367 (2021) doi:10.1101/2021.11.02.466367.

44. Hoikkala, V. et al. Cooperation between Different CRISPR-Cas Types Enables Adaptation in an RNA-Targeting System. MBio 12, (2021).

45. Mohanraju, P. et al. Alternative functions of CRISPR-Cas systems in the evolutionary arms race. Nat. Rev. Microbiol. 20, 351–364 (2022).

46. Landsberger, M. et al. Anti-CRISPR Phages Cooperate to Overcome CRISPR-Cas Immunity. Cell 174, 908–916.e12 (2018).

47. Hussain, F. A. et al. Rapid evolutionary turnover of mobile genetic elements drives bacterial resistance to phages. Science 374, 488–492 (2021).

48. Piel, D. et al. Phage-host coevolution in natural populations. Nat Microbiol 7, 1075–1086 (2022).

49. Doron, S.et al. Systematic discovery of antiphage defense systems in the microbial pangenome. Science 359, (2018).

50. Cohen, D. et al. Cyclic GMP-AMP signalling protects bacteria against viral infection. Nature 574, 691–695 (2019).

51. Birkholz, N. & Fineran, P. C. Turning down the (C)BASS: Phage-encoded inhibitors jam bacterial immune signaling. Molecular cell vol. 82 2185–2187 (2022).

52. Hobbs, S. J. et al. Phage anti-CBASS and anti-Pycsar nucleases subvert bacterial immunity. Nature 605, 522–526 (2022).

53. Gautreau, G. et al. PPanGGOLiN: Depicting microbial diversity via a partitioned pangenome graph. PLoS Comput. Biol. 16, e1007732 (2020).

54. Schwengers, O. et al. Bakta: rapid and standardized annotation of bacterial genomes via alignment-free sequence identification. Microb Genom 7, (2021).

55. Payne, L. J. et al. Identification and classification of antiviral defence systems in bacteria and archaea with PADLOC reveals new system types. Nucleic Acids Res. 49, 10868–10878 (2021).

56. Bernheim, A. & Sorek, R. The pan-immune system of bacteria: antiviral defence as a community resource. Nat. Rev. Microbiol. 18, 113–119 (2020).

57. Califano, J. V. et al. Characterization of Porphyromonas gingivalis insertion sequence-like element ISPg5. Infect. Immun. 68, 5247–5253 (2000).

58. Simpson Waltena et al. Transposition of the Endogenous Insertion Sequence Element IS1126 Modulates Gingipain Expression inPorphyromonas gingivalis. Infect. Immun. 67, 5012–5020 (1999).

59. Zhao, S. et al. Adaptive Evolution within Gut Microbiomes of Healthy People. Cell Host Microbe 25, 656–667.e8 (2019).

60. Silpe, J. E., Duddy, O. P., Hussain, F. A., Forsberg, K. J. & Bassler, B. L. Small protein modules dictate prophage fates during polylysogeny. bioRxiv 2022.09.16.508337 (2022) doi:10.1101/2022.09.16.508337.

61. Selva, L. et al. Killing niche competitors by remote-control bacteriophage induction. Proc. Natl. Acad. Sci. U. S. A. 106, 1234–1238 (2009).

62. Leke, N., Grenier, D., Goldner, M. & Mayrand, D. Effects of hydrogen peroxide on growth and selected properties of Porphyromonas gingivalis. FEMS Microbiol. Lett. 174, 347–353 (1999).

63. Ho, M.-H., Chen, C.-H., Goodwin, J. S., Wang, B.-Y. & Xie, H. Functional Advantages of Porphyromonas gingivalis Vesicles. PLoS One 10, e0123448 (2015).

64. Guerin, E. et al. Isolation and characterisation of ΦcrAss002, a crAss-like phage from the human gut that infects Bacteroides xylanisolvens. Microbiome 9, 89 (2021).

65. Owen, S. V. et al. A window into lysogeny: revealing temperate phage biology with transcriptomics. Microb Genom 6, (2020).

66. Waterbury, J. B. & Valois, F. W. Resistance to co-occurring phages enables marine synechococcus communities to coexist with cyanophages abundant in seawater. Appl. Environ. Microbiol. 59, 3393–3399 (1993).

67. Herelle, Felix d’ & Smith, George Hathorn. The bacteriophage, its rôle in immunity. 298 (Baltimore, Williams & Wilkins company, 1922).

68. Barr, J. J. et al. Bacteriophage adhering to mucus provide a non-host-derived immunity. Proc. Natl. Acad. Sci. U. S. A. 110, 10771–10776 (2013).

69. Genco, R. J. et al. The Subgingival Microbiome Relationship to Periodontal Disease in Older Women. J. Dent. Res. 98, 975–984 (2019).

70. Preus, H. R., Olsen, I. & Gjermo, P. Bacteriophage infection--a possible mechanism for increased virulence of bacteria associated with rapidly destructive periodontitis. Acta Odontol. Scand. 45, 49–54 (1987).

71. Camargo, A. P. et al. IMG/VR v4: an expanded database of uncultivated virus genomes within a framework of extensive functional, taxonomic, and ecological metadata. Nucleic Acids Res. (2022) doi:10.1093/nar/gkac1037.

72. Mark Welch, J. L., Rossetti, B. J., Rieken, C. W., Dewhirst, F. E. & Borisy, G. G. Biogeography of a human oral microbiome at the micron scale. Proc. Natl. Acad. Sci. U. S.A. 113, E791–800 (2016).

73. Zhang, M., Whiteley, M. & Lewin, G. R. Polymicrobial Interactions of Oral Microbiota: a Historical Review and Current Perspective. MBio 13, e0023522 (2022).

74. Meier-Kolthoff, J. P. & Göker, M. VICTOR: genome-based phylogeny and classification of prokaryotic viruses. Bioinformatics 33, 3396–3404 (2017).

75. Gilchrist, C. L. M. & Chooi, Y.-H. Clinker & clustermap.js: Automatic generation of gene cluster comparison figures. Bioinformatics (2021) doi:10.1093/bioinformatics/btab007.

76. Langmead, B. Aligning short sequencing reads with Bowtie. Curr. Protoc. Bioinformatics Chapter 11, Unit 11.7 (2010).

77. Guo, J. et al. VirSorter2: a multi-classifier, expert-guided approach to detect diverse DNA and RNA viruses. Microbiome 9, 37 (2021).

78. Nayfach, S. et al. CheckV assesses the quality and completeness of metagenome-assembled viral genomes. Nat. Biotechnol. 39, 578–585 (2021).

79. Tisza, M. J., Belford, A. K., Domínguez-Huerta, G., Bolduc, B. & Buck, C. B. Cenote-Taker 2 democratizes virus discovery and sequence annotation. Virus Evol 7, veaa100 (2021).

80. Sirén, K. et al. Rapid discovery of novel prophages using biological feature engineering and machine learning. NAR Genom Bioinform 3, lqaa109 (2021).

81. Kieft, K., Zhou, Z. & Anantharaman, K. VIBRANT: automated recovery, annotation and curation of microbial viruses, and evaluation of viral community function from genomic sequences. Microbiome 8, 90 (2020).

82. Roux, S. et al. Cryptic inoviruses revealed as pervasive in bacteria and archaea across Earth’s biomes. Nat Microbiol 4, 1895–1906 (2019).

83. Wyss, C. Growth of Porphyromonas gingivalis, Treponema denticola, T. pectinovorum, T. socranskii, and T. vincentii in a chemically defined medium. J. Clin. Microbiol. 30, 2225–2229 (1992).

84. Murugkar, P. et al. Identification of a growth factor required for culturing specific fastidious oral bacteria. J. Oral Microbiol. 15, 2143651 (2023).

85. Chen, S., Zhou, Y., Chen, Y. & Gu, J. fastp: an ultra-fast all-in-one FASTQ preprocessor. Bioinformatics 34, i884–i890 (2018).

86. Wick, R. Filtlong: quality filtering tool for long reads. (Github).

87. Wick, R. Porechop: adapter trimmer for Oxford Nanopore reads. (Github).

88. Wick, R. R., Judd, L. M., Gorrie, C. L. & Holt, K. E. Unicycler: Resolving bacterial genome assemblies from short and long sequencing reads. PLoS Comput. Biol. 13, e1005595 (2017).

89. Wick, R. R. & Holt, K. E. Polypolish: Short-read polishing of long-read bacterial genome assemblies. PLoS Comput. Biol. 18, e1009802 (2022).

90. Li, H. bwa: Burrow-Wheeler Aligner for short-read alignment (see minimap2 for long-read alignment). (Github).

91. Zimin, A. V. et al. The MaSuRCA genome assembler. Bioinformatics 29, 2669–2677 (2013).

92. Zimin, A. V. & Salzberg, S. L. The genome polishing tool POLCA makes fast and accurate corrections in genome assemblies. PLoS Comput. Biol. 16, e1007981 (2020).

93. Jakočiūnė, D. & Moodley, A. A Rapid Bacteriophage DNA Extraction Method. Methods Protoc 1, (2018).

94. Faust, G. G. & Hall, I. M. SAMBLASTER: fast duplicate marking and structural variant read extraction. Bioinformatics 30, 2503–2505 (2014).

95. Quinlan, A. R. & Hall, I. M. BEDTools: a flexible suite of utilities for comparing genomic features. Bioinformatics 26, 841–842 (2010).

96. Jain, C., Rodriguez-R, L. M., Phillippy, A. M., Konstantinidis, K. T. & Aluru, S. High throughput ANI analysis of 90K prokaryotic genomes reveals clear species boundaries. Nat. Commun. 9, 5114 (2018).

97. Ondov, B. D. et al. Mash: fast genome and metagenome distance estimation using MinHash. Genome Biol. 17, 132 (2016).

98. Han, M. V. & Zmasek, C. M. phyloXML: XML for evolutionary biology and comparative genomics. BMC Bioinformatics 10, 356 (2009).

99. Yutin, N., Puigbò, P., Koonin, E. V. & Wolf, Y. I. Phylogenomics of prokaryotic ribosomal proteins. PLoS One 7, e36972 (2012).

100. Milicevic, O., Repac, J., Bozic, B., Djordjevic, M. & Djordjevic, M. A Simple Criterion for Inferring CRISPR Array Direction. Front. Microbiol. 10, 2054 (2019).

101. Silas, S. et al. Type III CRISPR-Cas systems can provide redundancy to counteract viral escape from type I systems. Elife 6, (2017).

102. Vink, J. N. A., Baijens, J. H. L. & Brouns, S. J. J. PAM-repeat associations and spacer selection preferences in single and co-occurring CRISPR-Cas systems. Genome Biol. 22, 281 (2021).

103. Hyatt, D. et al. Prodigal: prokaryotic gene recognition and translation initiation site identification. BMC Bioinformatics 11, 119 (2010).

104. McNair, K., Zhou, C., Dinsdale, E. A., Souza, B. & Edwards, R. A. PHANOTATE: a novel approach to gene identification in phage genomes. Bioinformatics 35, 4537–4542 (2019).

105. Steinegger, M. & Söding, J. MMseqs2 enables sensitive protein sequence searching for the analysis of massive data sets. Nat. Biotechnol. 35, 1026–1028 (2017).

106. Terzian, P. et al. PHROG: families of prokaryotic virus proteins clustered using remote homology. NAR Genom Bioinform 3, lqab067 (2021).

107. Remmert, M., Biegert, A., Hauser, A. & Söding, J. HHblits: lightning-fast iterative protein sequence searching by HMM-HMM alignment. Nat. Methods 9, 173–175 (2011).

108. Huerta-Cepas, J. et al. eggNOG 5.0: a hierarchical, functionally and phylogenetically annotated orthology resource based on 5090 organisms and 2502 viruses. Nucleic Acids Res. 47, D309–D314 (2019).

109. Cantalapiedra, C. P., Hernández-Plaza, A., Letunic, I., Bork, P. & Huerta-Cepas, J. eggNOG-mapper v2: Functional Annotation, Orthology Assignments, and Domain Prediction at the Metagenomic Scale. Mol. Biol. Evol. 38, 5825–5829 (2021).

110. Marchler-Bauer, A. & Bryant, S. H. CD-Search: protein domain annotations on the fly. Nucleic Acids Res. 32, W327–31 (2004).

111. Marchler-Bauer, A. et al. CDD: a Conserved Domain Database for the functional annotation of proteins. Nucleic Acids Res. 39, D225–9 (2011).

112. Marchler-Bauer, A. et al. CDD: NCBI’s conserved domain database. Nucleic Acids Res. 43, D222–6 (2015).

113. Marchler-Bauer, A. et al. CDD/SPARCLE: functional classification of proteins via subfamily domain architectures. Nucleic Acids Res. 45, D200–D203 (2017).

114. Lu, S. et al. CDD/SPARCLE: the conserved domain database in 2020. Nucleic Acids Res. 48, D265–D268 (2020).

115. Kelley, L. A., Mezulis, S., Yates, C. M., Wass, M. N. & Sternberg, M. J. E. The Phyre2 web portal for protein modeling, prediction and analysis. Nat. Protoc. 10, 845–858 (2015).

116. Zimmermann, L. et al. A Completely Reimplemented MPI Bioinformatics Toolkit with a New HHpred Server at its Core. J. Mol. Biol. 430, 2237–2243 (2018).

117. Teufel, F. et al. SignalP 6.0 predicts all five types of signal peptides using protein language models. Nat. Biotechnol. 40, 1023–1025 (2022).

118. Potter, S. C. et al. HMMER web server: 2018 update. Nucleic Acids Res. 46, W200–W204 (2018).

119. Ramsey, J. et al. Galaxy and Apollo as a biologist-friendly interface for high-quality cooperative phage genome annotation. PLoS Comput. Biol. 16, e1008214 (2020).

120. Lopes, A., Tavares, P., Petit, M.-A., Guérois, R. & Zinn-Justin, S. Automated classification of tailed bacteriophages according to their neck organization. BMC Genomics 15, 1027 (2014).

121. Nishimura, Y. et al. ViPTree: the viral proteomic tree server. Bioinformatics 33, 2379–2380 (2017).

122. Moraru, C., Varsani, A. & Kropinski, A. M. VIRIDIC—A novel tool to calculate the intergenomic similarities of prokaryote-infecting viruses. Viruses (2020).

123. Letunic, I. & Bork, P. Interactive Tree Of Life (iTOL) v5: an online tool for phylogenetic tree display and annotation. Nucleic Acids Res. 49, W293–W296 (2021).

124. Duran-Pinedo, A. E. et al. Community-wide transcriptome of the oral microbiome in subjects with and without periodontitis. ISME J. 8, 1659–1672 (2014).

125. Chen, T. et al. The Human Oral Microbiome Database: a web accessible resource for investigating oral microbe taxonomic and genomic information. Database 2010, baq013 (2010).

